# Purine permease 5 contributes to riboflavin distribution in *Arabidopsis* reproductive organs

**DOI:** 10.64898/2026.02.25.708111

**Authors:** Rui Shibata, Hikari Kuwata, Takuto Sugimoto, Madoka Kikuchi, Takahiro Ishikawa, Takahisa Ogawa

## Abstract

Riboflavin (vitamin B□; RF) and its derivatives flavin mononucleotide (FMN) and flavin adenine dinucleotide (FAD) are indispensable cofactors for redox reactions in plants. While higher plants possess a conserved pathway for *de novo* riboflavin biosynthesis, how flavins are transported and spatially distributed between tissues remains unresolved. In particular, the molecular identity of plasma membrane transporters involved in flavin transport in plants remains largely unknown. Here, we identify *Arabidopsis* purine permease 5 (*AtPUP5*) as a plasma membrane–localized protein associated with riboflavin transport and its distribution in plants. Using a riboflavin-auxotrophic yeast mutant, we show that *AtPUP5* enhances intracellular accumulation of RF and, to a lesser extent, FMN, whereas FAD accumulation is inefficient under the conditions tested. In planta, *AtPUP5* overexpression increases RF accumulation following external application. In contrast, loss-of-function mutants do not display defects in bulk RF uptake at the whole-plant level, indicating that *AtPUP5* is not essential for global riboflavin acquisition. Notably, *AtPUP5* deficiency results in RF overaccumulation in reproductive organs, including inflorescences, siliques, and seeds, irrespective of external RF supply. This organ-specific phenotype is fully suppressed by genetic complementation and coincides spatially with strong *AtPUP5* promoter activity in reproductive tissues. These findings suggest that AtPUP5 contributes to the regulation of localized riboflavin distribution in reproductive tissues. Together, our study provides a molecular framework for plasma membrane–mediated regulation of riboflavin distribution in plants and highlights spatial regulation as an important component of flavin homeostasis.

## Introduction

Riboflavin (vitamin B□; RF) is a water-soluble vitamin that predominantly occurs in its phosphorylated and adenylated forms, flavin mononucleotide (FMN) and flavin adenine dinucleotide (FAD), as cofactors of flavoproteins. These flavin cofactors reversibly interconvert among oxidized, semiquinone, and reduced states, thereby enabling a wide spectrum of redox reactions through their distinctive chemical properties (Powers, 2003). FMN and FAD serve as indispensable cofactors for numerous oxidoreductases and are therefore essential for primary metabolism across all living organisms.

In higher plants, flavins participate not only in central metabolic processes such as photosynthesis, respiration, and fatty acid metabolism, but also in the biosynthesis of vitamins B□, B□□, C, and folate. Furthermore, they act as cofactors for photoreceptors and DNA repair enzymes (Bacher et al., 2000; Roje, 2007). The essentiality of flavin metabolism for plant development is evident from the embryonic lethality of mutants lacking *PyrR* and the bleaching phenotype caused by reduced expression of *AtRIBA1* within the RIBA family (Hiltunen et al., 2012; Hasnain et al., 2013). RF has also been implicated as a signaling molecule involved in hormone responses (Dong & Beer, 2000; Boubakri et al., 2016).

Plants, bacteria, and fungi are capable of *de novo* RF biosynthesis *via* a pathway that is highly conserved across species. The pathway begins with one GTP molecule and two ribulose-5-phosphate molecules. GTP is converted through a four-step enzymatic sequence into 5-amino-6-ribitylamino-2,4(1H,3H)pyrimidinedione (ARP) (Herz et al., 2000; Hedtke et al., 2012; Hasnain et al., 2013; Sa et al., 2016), whereas ribulose-5-phosphate is transformed into 3,4-dihydroxy-2-butanone-4-phosphate by the bifunctional enzyme encoded by *AtRibA1* (Herz et al., 2000). These intermediates subsequently condense in a lumazine synthase–catalyzed reaction (Jordan et al., 1999; Xiao et al., 2004), and the resulting lumazine is converted into riboflavin by RF synthase (Fischer et al., 2005). RF is then further converted to FMN and FAD through ATP-dependent phosphorylation by RF kinase and adenylation by FAD synthase, respectively (Roje, 2007; Lynch & Roje, 2022).

Although RF biosynthesis in plants is thought to occur primarily in plastids (Gerdes et al., 2012; Eggers et al., 2021), enzymatic activities involved in FMN and FAD metabolism, including RF kinase, are also present in the cytosol and mitochondria (Sandoval & Roje, 2005; Giancaspero et al., 2009; Rawat et al., 2011). These observations indicate that flavins must be transported and dynamically distributed across multiple subcellular compartments and tissues. However, the molecular mechanisms underlying flavin transport at all hierarchical levels—including inter-organelle, intercellular, and inter-organ transport—remain largely unknown.

In contrast to plants, mammals lack the ability to synthesize RF and therefore rely entirely on extracellular uptake. In bacteria, several RF transporters have been identified, including YpaA in *Bacillus subtilis*, PnuX in *Corynebacterium glutamicum*, and RibU in *Lactococcus lactis* (Burgess et al., 2006; Duurkens et al., 2007; Vogl et al., 2007). Many of these bacterial genes are transcriptionally regulated by FMN riboswitches, demonstrating tight coupling between flavin metabolism and transport at the level of gene expression (Winkler et al., 2002; Burgess et al., 2006; Vogl et al., 2007).

In eukaryotes, flavin transport mechanisms have also been molecularly characterized. The yeast *Saccharomyces cerevisiae* employs Mch5p as a facilitated-diffusion RF transporter at the plasma membrane (Reihl & Stolz, 2005), and the mitochondrial carrier Flx1p mediates FAD exchange across the inner mitochondrial membrane (Bafunno et al., 2004). In mammals, three highly selective RF transporters, RFVT1-3, encoded by *SLC52A1-A3*, have been characterized, and mutations in *SLC52A2* and *SLC52A3* underlie several hereditary neuropathies (Jin & Yonezawa, 2022).

Despite these advances in other organisms, the molecular identity of flavin transporters in higher plants remains largely unknown. Although carrier-mediated RF uptake has been demonstrated in mitochondria isolated from tobacco BY-2 cells (Giancaspero et al., 2009), and although RF secretion from roots under iron deficiency is well documented (Gheshlaghi et al., 2021), no corresponding transporters have been identified. Moreover, little is known about the movement of RF, FMN, or FAD between plastids and mitochondria, between cells or organs, or between plants and their external environment. Thus, while physiological evidence suggests flavin mobility, its molecular basis in plants remains undefined. Such spatial regulation of flavin distribution is likely critical for coordinating metabolic activity across tissues and organelles. As a result, the regulatory framework governing flavin homeostasis in plants remains a major unresolved question in plant metabolism.

Given their demonstrated capacity to transport structurally diverse heterocyclic compounds—including adenine, pyridoxine, cytokinin, caffeine, nicotine, and various alkaloids—members of the purine permease (PUP) family emerge as compelling candidates for flavin transport in plants (Gillissen et al., 2000; Bürkle et al., 2003; Hildreth et al., 2011; Dastmalchi et al., 2019). Notably, RF shares key structural features with known PUP substrates, including a heterocyclic ring system and polar substituents, which raises the possibility that RF may also be accommodated by PUP-mediated transport mechanisms. Functional diversification within the PUP family has been documented across species, including nicotine transport by tobacco NUP1, cytokinin transport by rice OsPUP7, and adenine/caffeine transport in coffee and tea (Hildreth et al., 2011; Qi & Xiong, 2013; Kakegawa et al., 2019; Zhang et al., 2022). *Arabidopsis* contains 21 PUP family members, among which AtPUP1 and AtPUP2 transport adenine and pyridoxine, whereas AtPUP14 functions as a cytokinin transporter (Gillissen et al., 2000; Bürkle et al., 2003; Szydlowski et al., 2013; Zürcher et al., 2016). Recent large-scale functional analyses have further highlighted functional diversification within the PUP family. For example, a genome-wide multi-targeted CRISPR screen identified *PUP8*, together with *PUP7* and *PUP21*, as being involved in shoot growth and meristem-associated processes, implicating *PUP8* in cytokinin-related physiological functions rather than in primary metabolite transport (Hu et al., 2023). These findings support the idea that individual PUP members have evolved distinct substrate specificities and physiological roles.

Given this gap, understanding how flavins are transported and spatially distributed in plants remains an important challenge in plant metabolism. In this study, we identify *Arabidopsis* PURINE PERMEASE 5 (AtPUP5) as a plasma membrane protein associated with flavin transport. Through yeast complementation assays and genetic analyses in *Arabidopsis*, we provide evidence that AtPUP5 contributes to localized riboflavin distribution, particularly in reproductive tissues, revealing a previously unrecognized component of flavin homeostasis in plants.

## Results

### Functional complementation of the yeast rib5Δ mutant by Arabidopsis PUP family members

A previous study demonstrated that a *Saccharomyces cerevisiae* mutant lacking the *RIB5* gene (*rib5*Δ), which encodes the enzyme responsible for the final step of RF biosynthesis, is unable to synthesize RF endogenously and therefore requires exogenous RF for growth. Using this biosynthesis-deficient mutant, complementation assays previously identified the monocarboxylate transporter homolog Mch5p as an RF transporter that enhances RF uptake in *S. cerevisiae* (Reihl and Stolz, 2005).

To examine whether members of the *Arabidopsis* purine permease (AtPUP) family facilitate RF uptake, we employed the *S. cerevisiae rib5*Δ mutant as a heterologous expression system. Following the strategy described by Reihl and Stolz (2005), we generated a *rib5*Δ mutant strain (see Materials and Methods). As previously reported, the *rib5*Δ mutant exhibited growth only in medium supplemented with a high concentration of RF, consistent with its inability to synthesize RF (Figure S1).

Total RNA was extracted from the aerial parts of *Arabidopsis thaliana*, and cDNAs corresponding to 18 of the 21 *AtPUP* genes were amplified and cloned into the yeast expression vector pYES2. cDNAs for AtPUP13, AtPUP15, and AtPUP20 were not obtained from aerial tissue-derived RNA and were therefore excluded from subsequent yeast assays. The resulting constructs were transformed into the *rib5*Δ mutant and subjected to growth assays on uracil-deficient synthetic medium (SD/–Ura) and galactose-containing induction medium (SG/–Ura) supplemented with RF (0, 0.2, 2, or 20 mg L□¹).

Under low RF conditions, the empty vector control and most *AtPUP*-expressing strains failed to grow (Figure 1a). In contrast, strains expressing *AtPUP5* or *AtPUP8* displayed detectable growth recovery at these low RF concentrations, suggesting that *AtPUP5* and *AtPUP8* may enhance cellular uptake of exogenously supplied RF (Figure 1a).

**Figure 1.**
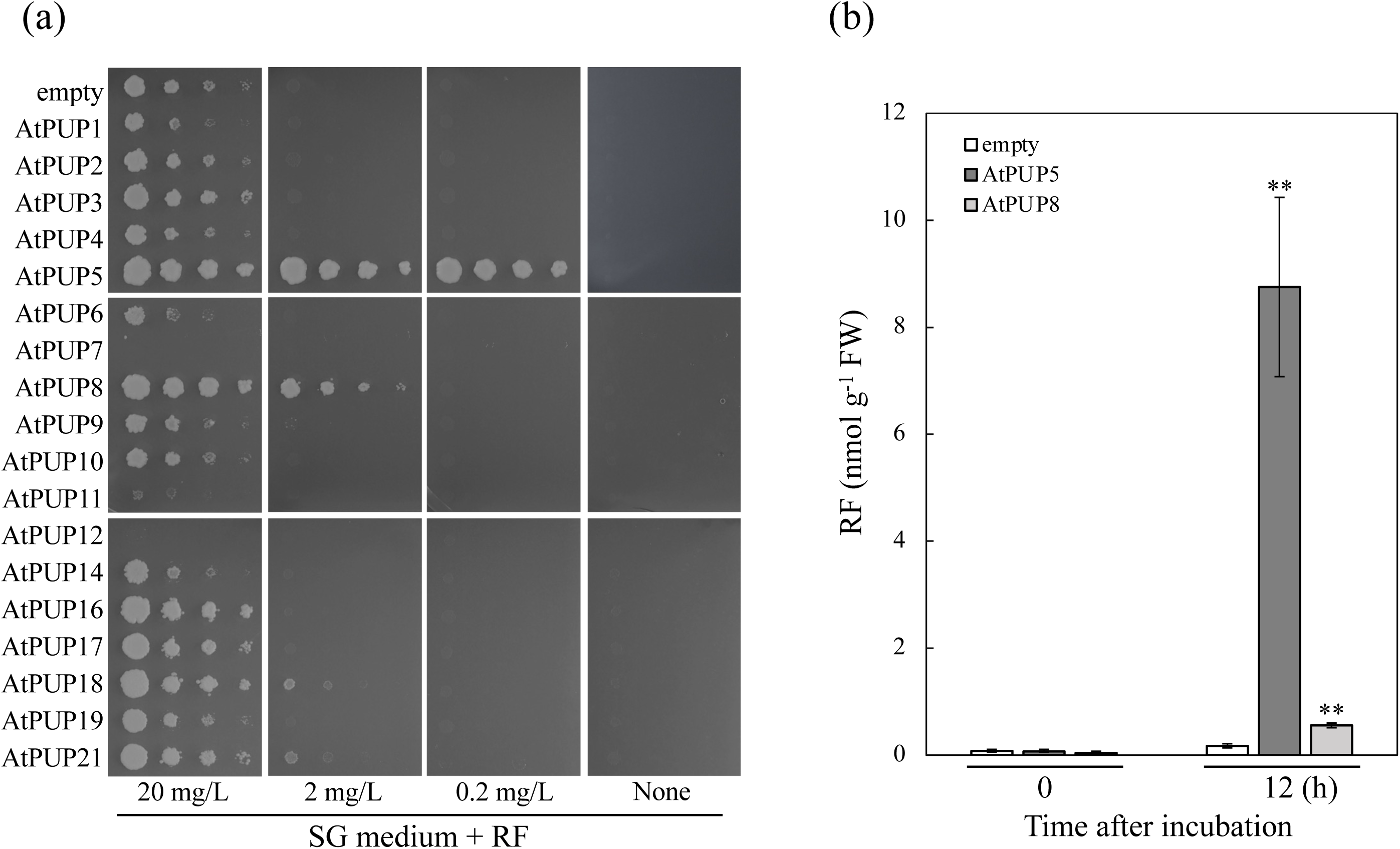
Functional complementation of the *Saccharomyces cerevisiae rib5*Δ mutant by *Arabidopsis* purine permeases. (a) A complementation assay was performed using the *S. cerevisiae rib5*Δ mutant, which lacks the gene encoding the final enzyme of RF synthesis. The mutant was transformed with an empty pYES2 vector (negative control) or pYES2 constructs harboring individual *A. thaliana* PUP genes. Serial dilutions of the transformed strains were spotted onto synthetic defined media lacking uracil and containing glucose (SD/−Ura) or galactose (SG/−Ura), supplemented with RF at the indicated concentrations (0, 0.2, 2, 20 mg L^-1^), and incubated at 30 °C for 72 h. (b) Intracellular RF levels were quantified in *rib5*Δ yeast cells expressing *AtPUP5* or *AtPUP8* after incubation for 12 h in medium supplemented with 50 µM RF. Yeast cultures were grown in three independent biological replicates, and RF was quantified from harvested cells of each replicate. Data are presented as means ± SE (n = 3). Asterisks indicate statistically significant differences compared with the empty vector control at 12 h (Student’s *t*-test; \*\**P* < 0.01).

To evaluate RF uptake activity more directly, intracellular RF levels were quantified in yeast strains expressing *AtPUP5* or *AtPUP8*. Twelve hours after RF addition, intracellular RF levels in the *AtPUP5*-expressing strain increased to approximately 100-fold relative to the empty vector control, whereas the *AtPUP8*-expressing strain showed only a ∼10-fold increase (Figure 1b). These results indicate that *AtPUP5* is associated with substantially stronger RF uptake activity than *AtPUP8* in yeast.

### Short-term time-course analysis of flavin uptake in yeast cells expressing *AtPUP5*

To further characterize time-dependent flavin uptake, intracellular flavin levels were quantified over a short time course. In the *AtPUP5*-expressing strain, intracellular RF levels increased in a time-dependent manner beginning within 2 min after RF addition, reaching approximately 100-fold relative to the empty vector control strain (Figure 2a). Concomitant with the increase in intracellular RF, intracellular FMN levels also increased approximately threefold, which may reflect intracellular conversion of imported RF to FMN. Similarly, when FMN was supplied, a time-dependent increase in intracellular FMN levels was observed, accompanied by an increase in intracellular RF levels (Figure 2b), suggesting interconversion between flavin forms within yeast cells. In contrast, FAD supplementation did not result in a time-dependent increase in intracellular FAD levels, and intracellular RF and FMN levels did not change significantly over time (Figure 2c).

**Figure 2.**
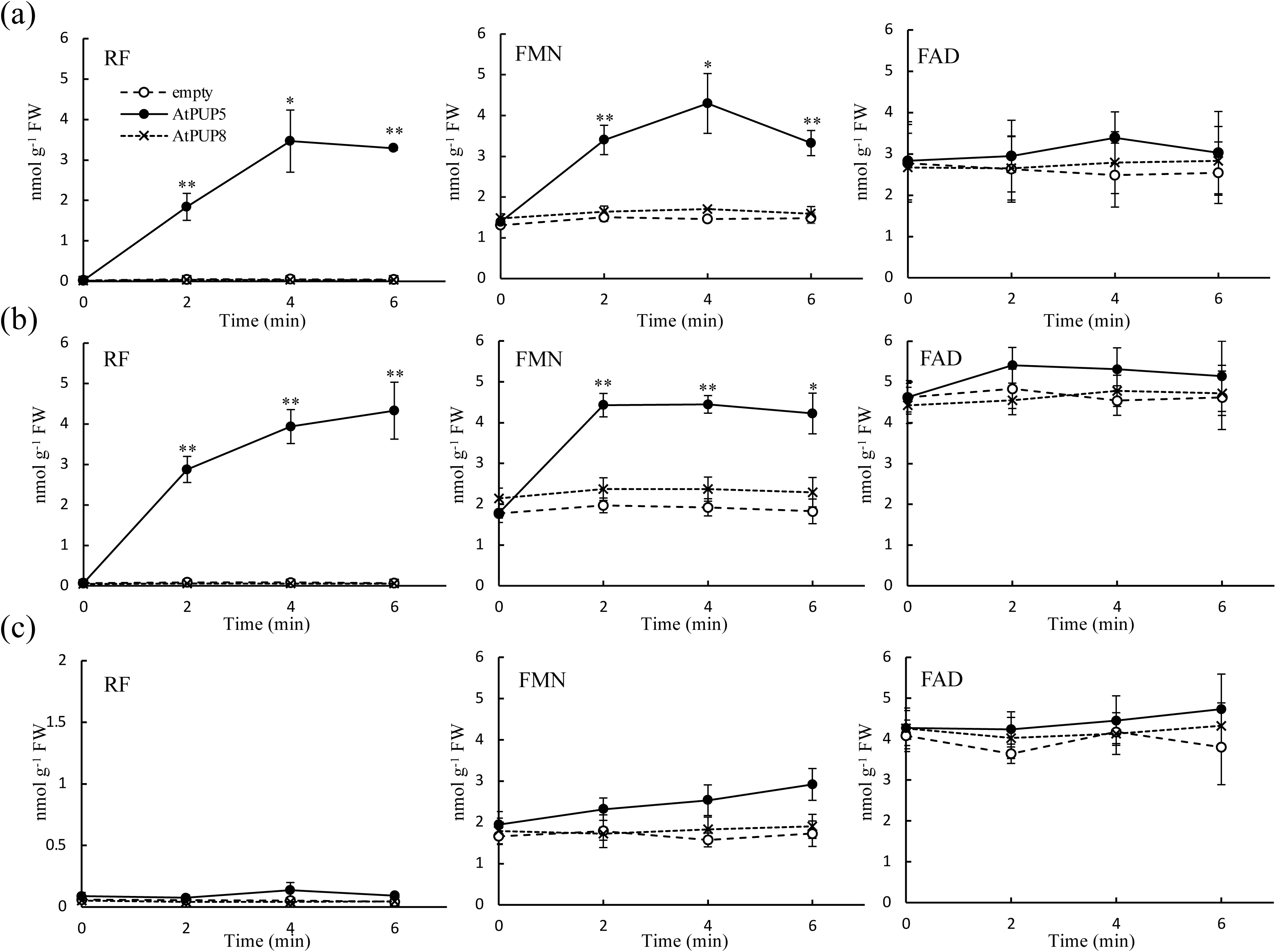
Short-term time-course analysis of flavin uptake in *S. cerevisiae rib5*Δ cells expressing AtPUP5 *or* AtPUP8. Time-course analyses of intracellular flavin levels were performed using the *rib5*Δ yeast cells expressing *AtPUP5* or *AtPUP8* following short-term incubation (2–6 min) with 50 µM (a) riboflavin (RF), (b) flavin mononucleotide (FMN), or (c) flavin adenine dinucleotide (FAD). Yeast cultures were grown in three independent biological replicates, and harvested cells from each replicate were analyzed. Data are presented as means ± SE (n = 3). Asterisks indicate statistically significant differences compared with the empty vector control at each time point (Student’s *t*-test; \**P* < 0.05 and \*\**P* < 0.01).

In the *AtPUP8*-expressing strain, no significant increase in intracellular flavin levels was detected during the short time course following the addition of RF, FMN, or FAD (Figure 2a–c). Together, these results suggest that *AtPUP5* enhances the uptake of RF and, to a lesser extent, FMN in yeast, whereas uptake of FAD appears to be substantially less efficient under the conditions tested. In contrast, *AtPUP8* did not show detectable flavin uptake during the short time course under the conditions tested. To place these functional differences in an evolutionary context, a phylogenetic analysis of purine permease (PUP) family members from *Arabidopsis thaliana* and *Oryza sativa* was performed (Figure S2). The resulting tree indicates that *AtPUP5* and *AtPUP8* belong to distinct clades, with *AtPUP5* clustering together with *OsPUP5*. Consistent with this phylogenetic relationship, expression of a rice PUP homolog closely related to *AtPUP5* (*OsPUP5*), originally described by Yamaki et al. (2017), also partially restored the growth of the *rib5*Δ mutant (Figure S3), supporting the possibility of functional conservation of flavin transport activity within the PUP5 subclade.

### AtPUP5, but not AtPUP8, enhances riboflavin accumulation in *Arabidopsis*

To examine whether AtPUP5 and AtPUP8 influence RF accumulation in planta, we generated *Arabidopsis* transgenic lines overexpressing *AtPUP5* or *AtPUP8*. Overexpression lines were produced by *Agrobacterium*-mediated transformation using constructs in which each cDNA was placed under the control of the Cauliflower mosaic virus 35S promoter. Quantitative RT–PCR analysis confirmed strong overexpression of the introduced genes, with transcript levels increased approximately 59-fold in OE-*AtPUP5* 2–6, 23-fold in OE-*AtPUP5* 4–1, and 100-fold in OE-*AtPUP8* 3–1 compared with the empty control (Figure S4).

Riboflavin accumulation following external RF application was evaluated using these overexpression lines. RF levels in the roots of the *AtPUP5*-overexpressing lines were significantly higher than those in the control, reaching approximately 13-fold and 9-fold relative to the empty vector control, respectively, 6 h after RF treatment (Figure 3). In contrast, RF levels in both shoots and roots of the *AtPUP8*-overexpressing line were comparable to those of the control (Figure 3). These results suggest that *AtPUP5*, but not *AtPUP8*, promotes RF accumulation in *Arabidopsis* following external application. Given this pronounced difference in RF accumulation, subsequent analyses primarily focused on *AtPUP5*.

**Figure 3.**
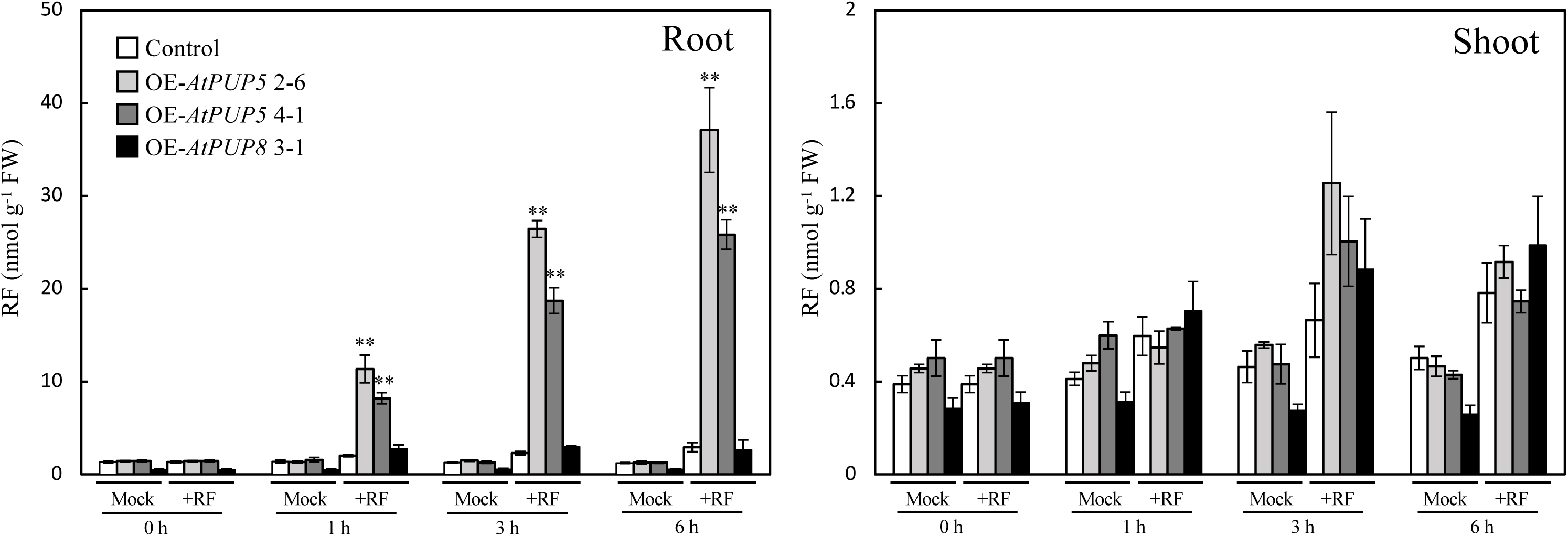
Riboflavin accumulation in *Arabidopsis* transgenic lines overexpressing *AtPUP5* or *AtPUP8* following external application. Control plants (wild-type plants transformed with an empty vector), two independent *AtPUP5*-overexpressing lines, and one independent *AtPUP8*-overexpressing line were analyzed. Plants were grown on half-strength Murashige and Skoog (1/2 MS) medium containing 1% (w/v) sucrose on cellophane sheets for 2 weeks under standard growth conditions. The seedlings grown on cellophane sheets were then transferred to filter paper soaked in liquid 1/2 MS medium supplemented with or without 50 µM riboflavin (RF) and incubated in the dark to prevent photodegradation of RF. After 1, 3, and 6 h of incubation, plants were harvested, separated into shoots and roots, thoroughly washed, and RF levels were quantified. Plants were grown independently in three biological replicates, and shoots and roots from each replicate were analyzed. Data are presented as means ± SE (n = 3). Asterisks indicate statistically significant differences compared with the empty vector control at each time point within each treatment (Student’s *t*-test; \*\**P* < 0.01).

To assess whether *AtPUP5* also affects the uptake of flavin coenzyme forms, FMN and FAD, these compounds were applied to the *AtPUP5*-overexpressing lines. FMN or FAD treatment did not result in detectable differences in intracellular FMN or FAD levels between the control and *AtPUP5*-overexpressing plants (Figure 4). However, following FMN treatment, RF levels in roots of the *AtPUP5*-overexpressing lines increased approximately 16-fold and 10-fold, respectively, relative to the empty vector control at 6 h (Figure 4a). Similarly, FAD treatment led to increases in root RF levels of approximately 16-fold and 7-fold, respectively, relative to the empty vector control (Figure 4b). These increases in RF levels may reflect metabolic conversion of exogenously supplied flavin coenzymes to riboflavin prior to or following uptake, rather than indicating efficient accumulation of FMN or FAD in planta.

**Figure 4.**
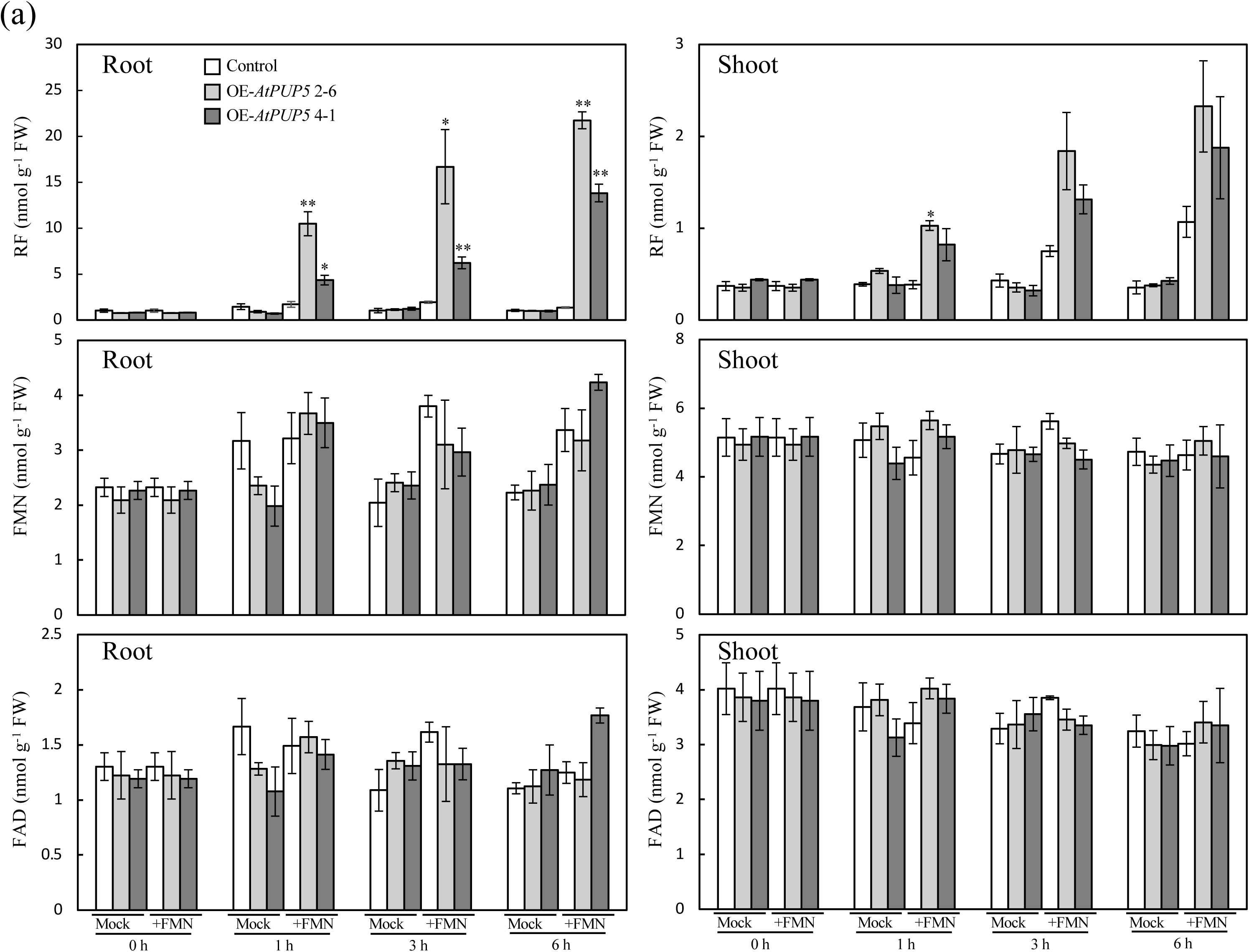
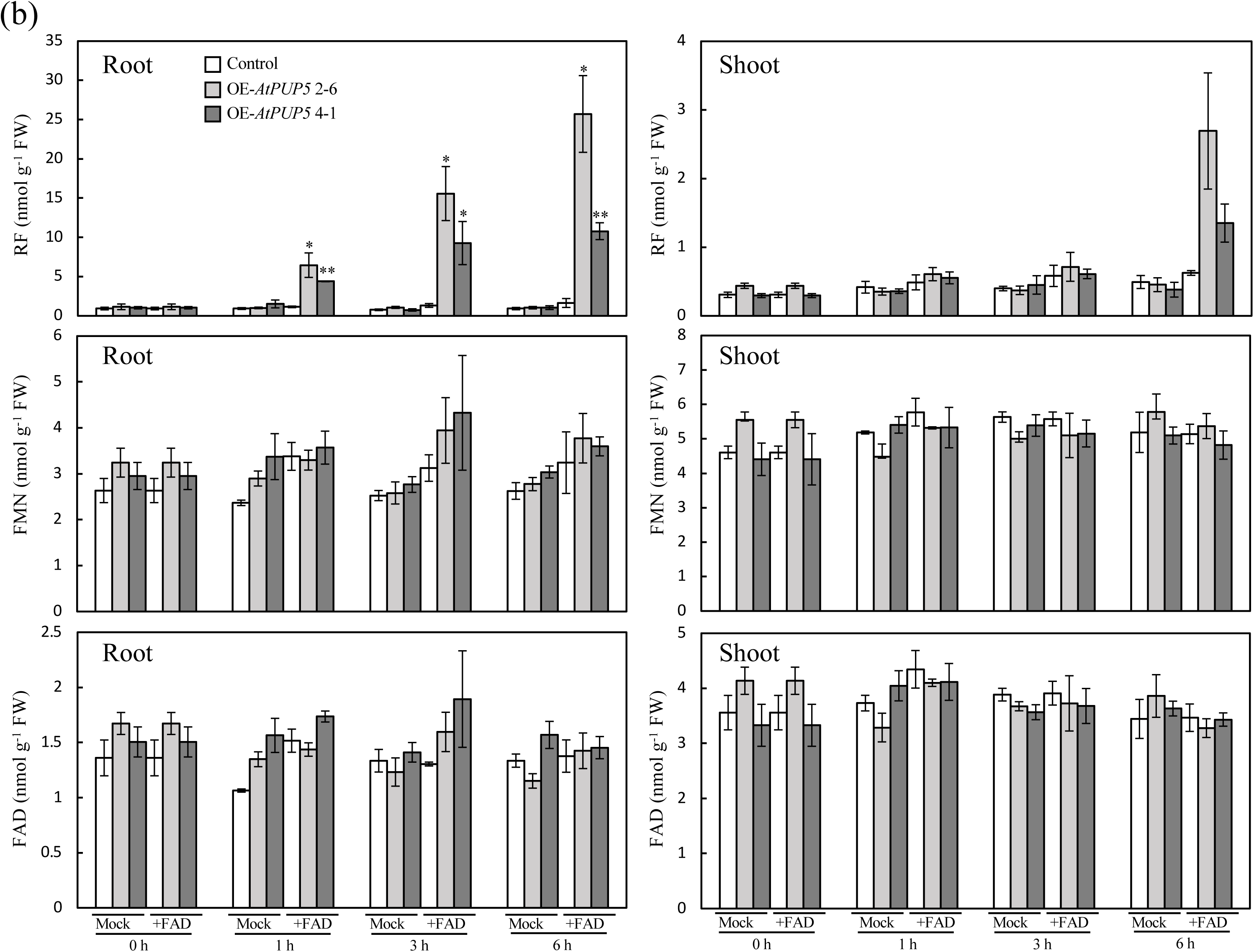
Riboflavin accumulation following application of flavin coenzymes in *Arabidopsis* plants overexpressing *AtPUP5*. Control plants (wild-type plants transformed with the empty vector) and two independent *AtPUP5*-overexpressing lines were analyzed. Plants were grown on half-strength Murashige and Skoog (1/2 MS) medium containing 1% (w/v) sucrose on cellophane sheets for 2 weeks under standard growth conditions. The seedlings grown on cellophane sheets were then transferred to filter paper soaked in liquid 1/2 MS medium supplemented with or without 50 µM (a) flavin mononucleotide (FMN) or (b) flavin adenine dinucleotide (FAD) and incubated in the dark to prevent photodegradation of flavins. After 1, 3, and 6 h of incubation, plants were harvested, separated into shoots and roots, thoroughly washed, and RF, FMN, and FAD levels were quantified. Plants were grown independently in three biological replicates, and shoots and roots from each replicate were analyzed. Data are presented as means ± SE (n = 3). Asterisks indicate statistically significant differences compared with the empty vector control at each time point within each treatment (Student’s *t*-test; \**P* < 0.05 and \*\**P* < 0.01).

### Loss of *AtPUP5* does not affect bulk riboflavin uptake but alters organ-specific riboflavin levels

To investigate the physiological roles of *AtPUP5* and *AtPUP8* in planta, T-DNA insertion knockout lines for *AtPUP5* and *AtPUP8* were isolated in *Arabidopsis*. Homozygous single knockout lines (*pup5* and *pup8*) as well as a double knockout line (*pup5/8*) were generated. Semi-quantitative RT–PCR analysis confirmed the absence of *AtPUP5* and/or *AtPUP8* transcripts in each respective mutant line (Figure 5a).

**Figure 5.**
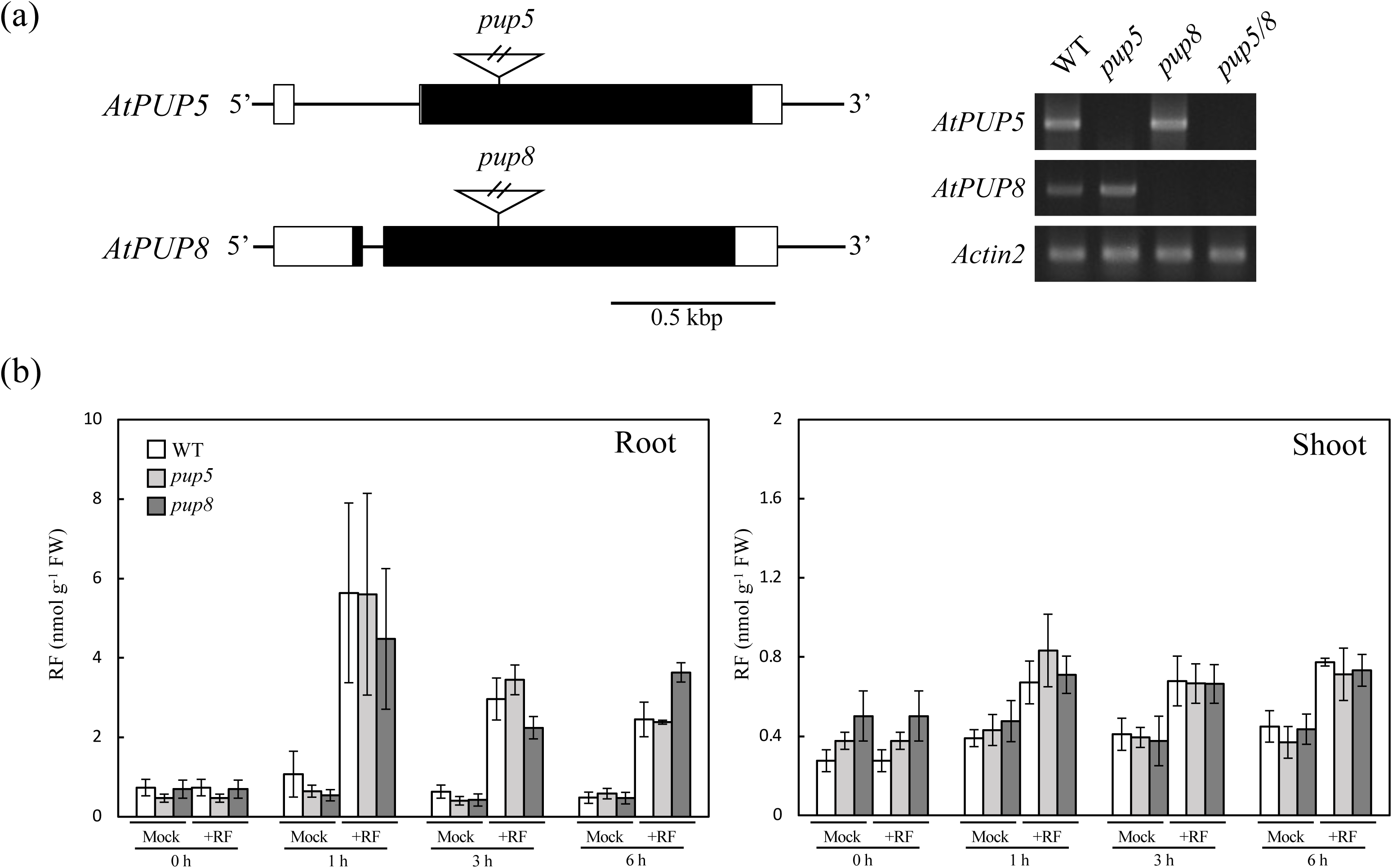
Effects of loss of *AtPUP5* and *AtPUP8* on riboflavin accumulation following external application. Wild-type plants and *AtPUP* knockout mutants (*pup5*, *pup8*, and *pup5/8*) were analyzed. Plants were grown on half-strength Murashige and Skoog (1/2 MS) medium supplemented with 1% (w/v) sucrose for 2 weeks under standard growth conditions. (a) T-DNA insertion sites in the *AtPUP5* and AtPUP8 genes of the *pup5* (CS868942) and *pup8* (SALK_137529) mutants, respectively, and semi-quantitative RT–PCR analysis of *AtPUP5* and *AtPUP8* transcripts in wild-type and mutant plants. T-DNA insertion sites are indicated by triangles, and black and white boxes represent exons and untranslated regions, respectively. (b) Quantification of riboflavin (RF) levels following external 50 µM RF application. Wild-type, *pup5*, and *pup8* plants were grown on 1/2 MS medium containing 1% (w/v) sucrose on cellophane sheets for 2 weeks under standard conditions. RF treatment and RF quantification were performed over a 1–6 h time course as described in Figure 3. Plants were grown independently in three biological replicates, and shoots and roots from each replicate were analyzed. Data are presented as means ± SE (n = 3).

Short-term RF uptake assays were performed to quantify RF levels over a 1–6 h time course following external application. RF levels in the shoots and roots of *pup5* and *pup8* mutants were comparable to those of the wild type at all examined time points, and no significant differences were detected among genotypes (Figure 5b).

To examine whether *AtPUP5* and *AtPUP8* contribute to RF uptake or accumulation over longer time scales, plants were grown in soil under normal conditions and irrigated with either RF-free or RF-supplemented water. Plant growth and development were evaluated, including measurements of leaf area and flowering time (days to bolting and number of leaves at bolting). No significant differences were observed between RF-treated and untreated plants, or between the wild type and the knockout lines, for any of these parameters (Figure S5).

Flavin levels were further quantified in various organs following long-term RF treatment. The extent of RF, FMN, and FAD accumulation in most organs did not differ significantly between the wild type and the *pup5*, *pup8*, or *pup5/8* mutants (Figure 6). These results suggest that loss of *AtPUP5* and *AtPUP8* does not markedly affect the overall accumulation of exogenously applied RF at the whole-plant level. Notably, we found that RF levels in the inflorescences and siliques of *pup5* and *pup5/8*, but not of *pup8* plants, were higher than those in the wild type, irrespective of RF treatment (Figure 6), suggesting an organ-specific alteration of riboflavin levels associated with loss of *AtPUP5*.

**Figure 6.**
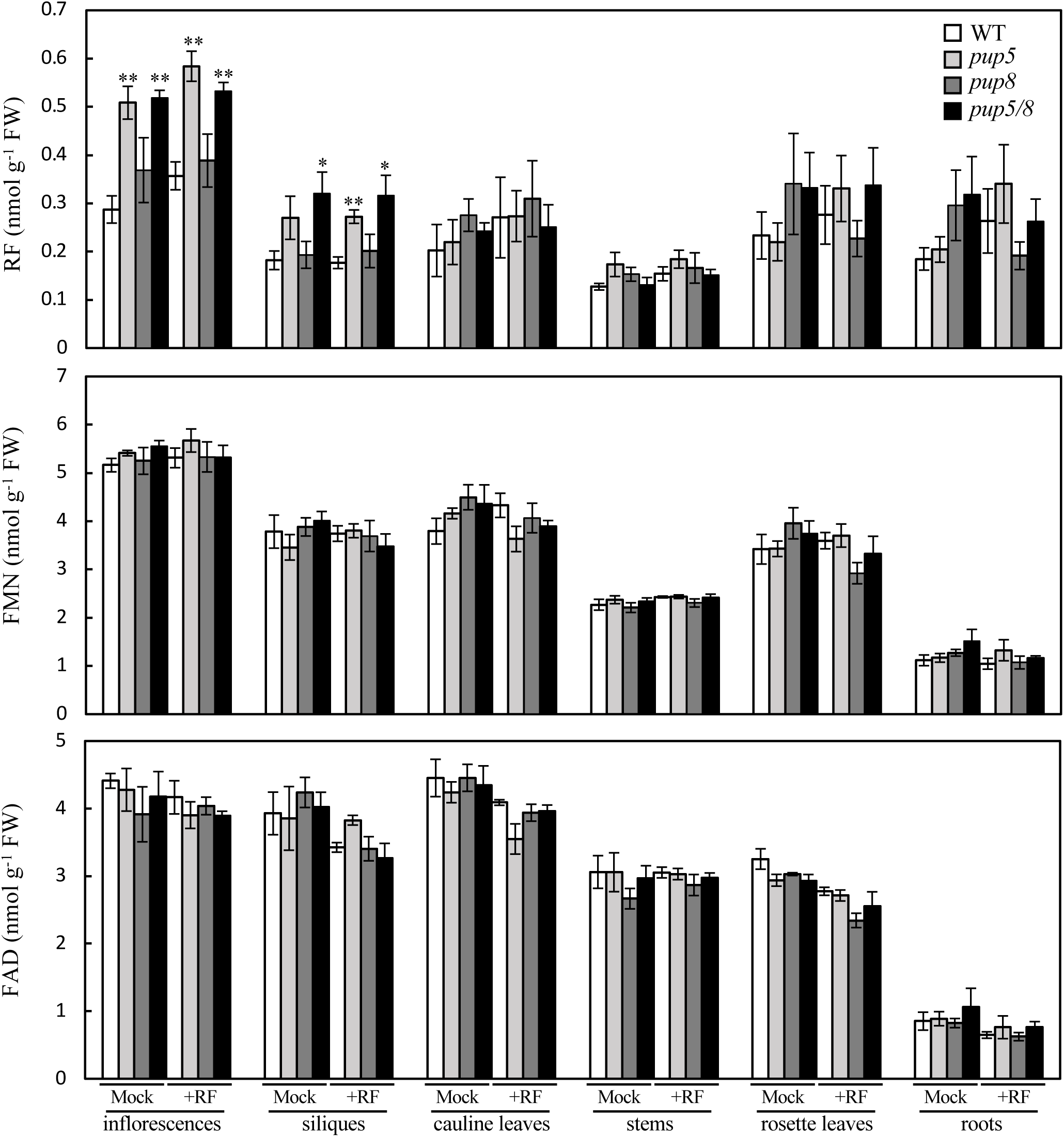
Effects of loss of *AtPUP5* and *AtPUP8* on flavin accumulation and organ-specific distribution following long-term riboflavin treatment. Wild-type plants and *AtPUP* knockout mutants (*pup5*, *pup8*, and *pup5/8*) were grown in soil for 40 days under standard growth conditions and irrigated with water (Mock) or water supplemented with 50 µM riboflavin (RF). At 40 days, organs from wild-type and mutant plants were harvested 4 h after the onset of illumination, and RF, FMN, and FAD levels were quantified. Plants were grown independently in three biological replicates, and organs from each replicate were analyzed. Data are presented as means ± SE (n = 3). Asterisks indicate statistically significant differences compared with the corresponding organ of wild-type plants within each treatment (Student’s *t*-test; \**P* < 0.05 and \*\**P* < 0.01).

### Loss of AtPUP5 leads to riboflavin overaccumulation in reproductive organs

To examine whether *AtPUP5* is involved in the distribution of flavin compounds among plant organs, flavin levels were quantified in various organs of the *pup5* mutant and complemented lines (*pup5*/*PUP5*). Semi-quantitative RT–PCR analysis confirmed that *AtPUP5* expression in the complemented lines was restored to levels comparable to or higher than those observed in the wild type (Figure S6).

Quantification of flavin compounds revealed that RF levels in inflorescences were significantly higher in *pup5* than in the wild type, whereas this increase was suppressed in the *pup5*/*PUP5* lines (Figure 7). In addition, RF levels in siliques and cauline leaves of *pup5* were also restored to the wild-type levels by complementation with *AtPUP5* (Figure 7a).

**Figure 7.**
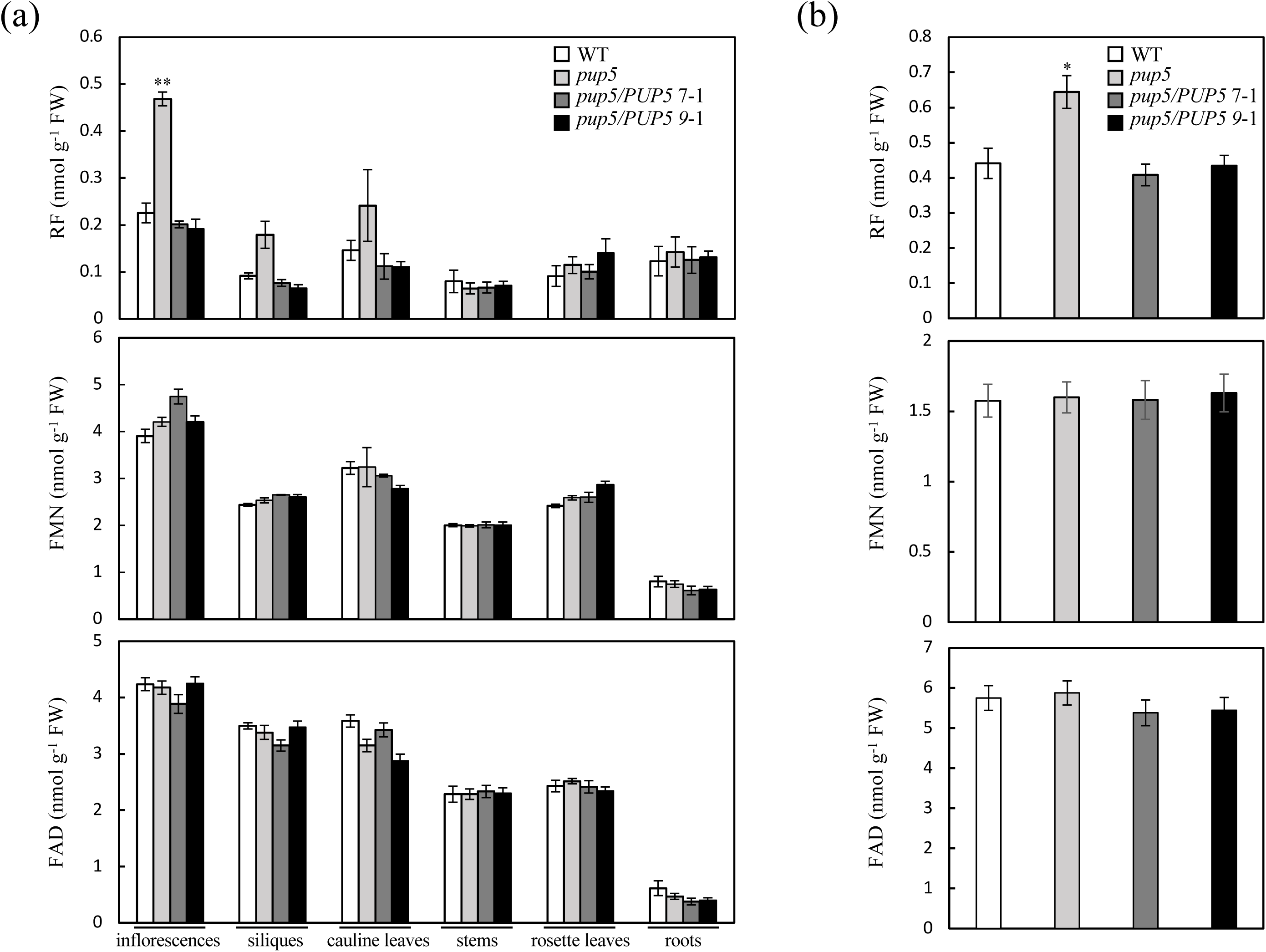
Organ-specific flavin accumulation in *pup5* mutant and *AtPUP5-*complemented lines. Wild-type plants, the *pup5* mutant, and two independent *AtPUP5*-complemented lines (*pup5*/*PUP5*) were analyzed. Plants were grown in soil under standard growth conditions. At 40 days, organs from wild-type, *pup5*, and complemented plants were harvested 4 h after the onset of illumination, and flavin compounds were quantified (a). Flavin levels in dry seeds harvested from wild-type, *pup5*, and complemented lines grown under standard growth conditions (b). Plants were grown independently in three biological replicates, and organs from each replicate were analyzed. Data are presented as means ± SE (n = 3). Asterisks indicate statistically significant differences compared with the corresponding wild-type organ (Student’s *t*-test; \**P* < 0.05 and \*\**P* < 0.01).

Because RF levels were elevated in reproductive tissues of *pup5*, flavin levels were further quantified in dry seeds. RF levels in seeds of *pup5* were significantly higher than those in the wild type (Figure 7b), and this increase was suppressed in the *pup5*/*PUP5* lines. Together, these results indicate that loss of *AtPUP5* results in organ-specific overaccumulation of RF, while overall flavin accumulation at the whole-plant level remains largely unchanged.

### AtPUP5 is expressed in reproductive tissues

To examine the tissue-specific expression pattern of *AtPUP5*, promoter–GUS reporter analysis was performed. A construct containing the *AtPUP5* promoter region (2,000 bp upstream of the translation start site) fused to the β-glucuronidase (GUS) reporter gene was introduced into wild-type *Arabidopsis* plants.

GUS staining of transgenic plants grown in soil for 40 days revealed strong promoter activity of *AtPUP5* in young rosette leaves and emerging aerial tissues (Figure 8a). The intensity of GUS staining in leaves gradually decreased as plants matured (Figure 8b). Prominent GUS activity was also detected in reproductive tissues, including inflorescences and siliques (Figure 8c). In siliques, GUS staining was strongest at early developmental stages and declined as siliques matured (Figure 8d).

**Figure 8.**
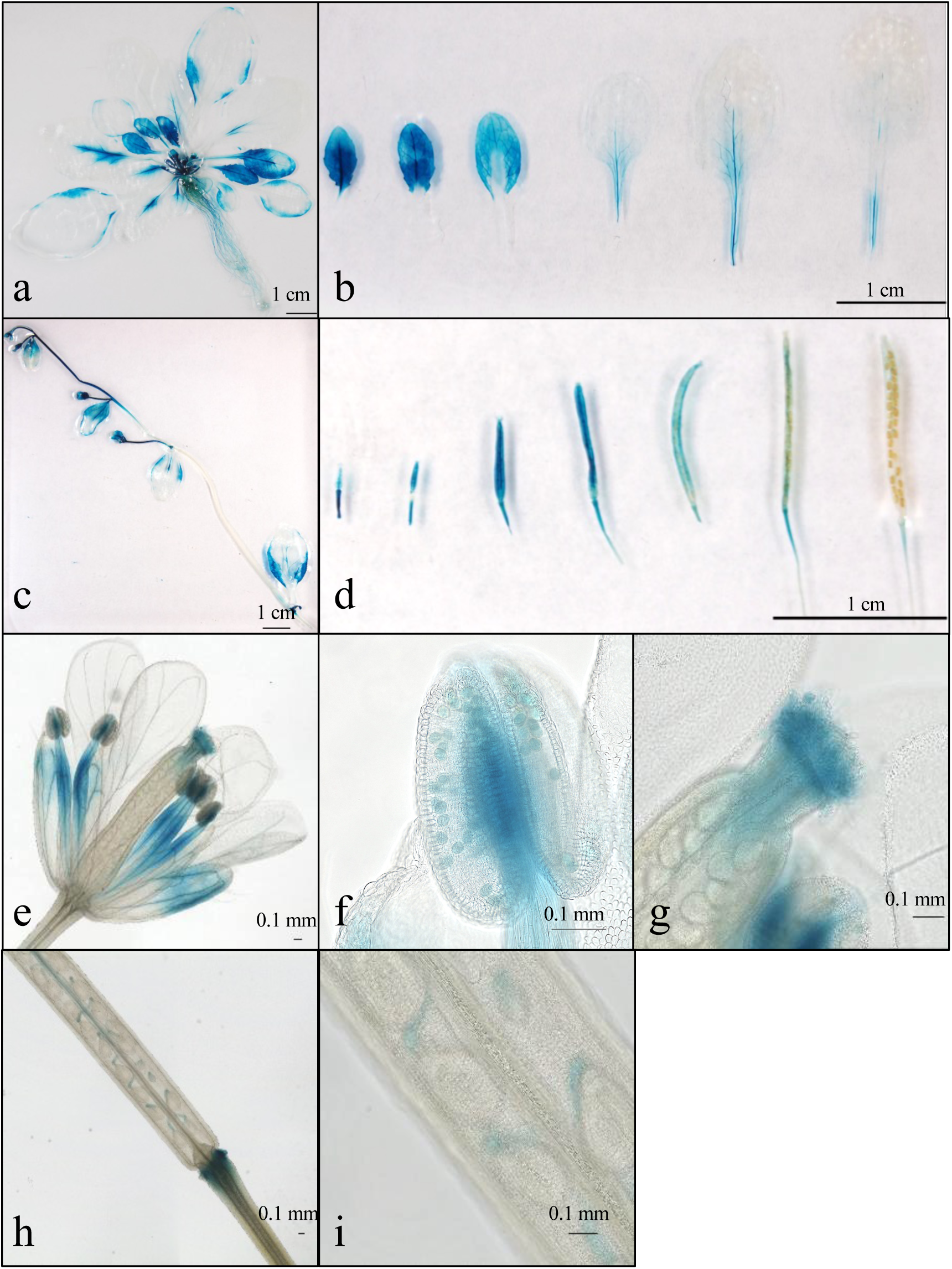
Tissue-specific expression pattern of *AtPUP5* revealed by promoter–GUS fusion analysis. Forty-day-old *Arabidopsis thaliana* transformed with a promoter–GUS fusion construct containing a 2.0-kb *AtPUP5* promoter region were stained with 5-bromo-4-chloro-3-indolyl β-D-glucuronide (X-Gluc) for 2 h, destained, and documented. Representative images show (a) whole plant, (b) rosette leaves, (c) stem including inflorescences and cauline leaves, and (d) siliques. Magnified views show (e) inflorescences, (f) anthers, (g) stigmas, and (h, i) siliques at different developmental stages.

Within inflorescences, *AtPUP5* promoter activity was particularly strong in specific reproductive structures, with intense staining observed at the tips of anthers and in stigmas (Figure 8e–i). This spatial expression pattern is consistent with the organ-specific RF accumulation observed in the *pup5* mutant. Quantitative RT–PCR analysis further confirmed that *AtPUP5* is preferentially expressed in reproductive tissues, including inflorescences and siliques (Figure S7).

### AtPUP5 localizes to the plasma membrane

To determine the subcellular localization of *AtPUP5* in *Arabidopsis*, a construct expressing an N-terminal GFP fusion of *AtPUP5* (nGFP–AtPUP5) was generated. The nGFP–AtPUP5 construct was co-introduced with a plasma membrane marker (PM–RFP) (Nishimura et al., 2016) into *Arabidopsis* leaf epidermal cells using particle bombardment. GFP and RFP fluorescence signals were subsequently observed by confocal laser scanning microscopy. The GFP fluorescence signal derived from nGFP–AtPUP5 overlapped extensively with the PM–RFP signal, indicating colocalization with the plasma membrane marker (Figure 9). These results suggest that *AtPUP5* localizes predominantly to the plasma membrane in plant cells.

**Figure 9.**
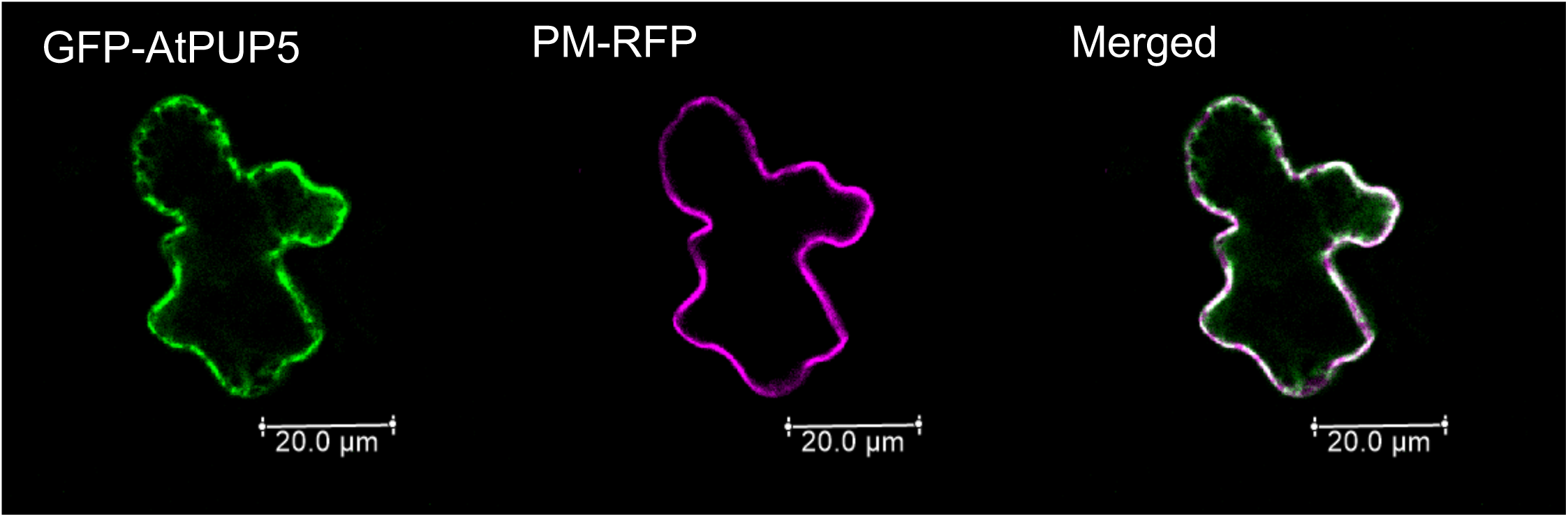
Plasma membrane localization of *AtPUP5* in plant cells. An N-terminal GFP fusion of *AtPUP5* (nGFP–AtPUP5; pGWB506/AtPUP5) was co-expressed with a plasma membrane–localized RFP marker (AtSYP132–RFP; pB5tdGW/AtSYP132) in *Arabidopsis thaliana* leaf epidermal cells by particle bombardment. Fluorescence signals derived from nGFP–AtPUP5 (left) and the plasma membrane marker RFP (center) were recorded by confocal laser scanning microscopy and merged (right).

## Discussion

In this study, we identify *AtPUP5* as a plasma membrane–localized protein associated with flavin transport and riboflavin distribution in higher plants, thereby addressing a long-standing gap in the understanding of flavin transport in plants. Our findings suggest that *AtPUP5* functions to regulate localized riboflavin distribution in *Arabidopsis thaliana*, providing molecular evidence that flavin transport contributes to organ- and tissue-specific riboflavin distribution in plants.

Analysis of RF uptake using *Arabidopsis* overexpression lines revealed that RF levels were significantly increased in *AtPUP5*-overexpressing plants, whereas no such increase was observed in *AtPUP8*-overexpressing plants (Figure 3). These results suggest that *AtPUP5*, but not *AtPUP8*, enhances RF accumulation in planta under the conditions examined. The absence of increased RF levels in the *AtPUP8* overexpression line suggests that RF is unlikely to be a major physiological substrate of *AtPUP8*, highlighting functional diversification within the PUP family. This interpretation is consistent with our yeast complementation and uptake assays, in which *AtPUP8* exhibited markedly lower growth rescue and flavin uptake activity than *AtPUP5* (Figures 1 and 2). A rice PUP homolog closely related to *AtPUP5* also partially restored growth of a flavin-auxotrophic yeast mutant (Figure S3), further suggesting that flavin-associated transport activity may be conserved within the *PUP5* subclade. Moreover, phylogenetic analysis indicates that *AtPUP5* and *AtPUP8* diverged early during evolution (Figure S2), which is consistent with the idea that these two transporters have acquired distinct physiological functions. Recent large-scale multi-targeted CRISPR analyses have further substantiated this functional divergence by demonstrating that *AtPUP8*, together with *PUP7* and *PUP21*, is involved in shoot growth and meristem-associated processes, implicating *AtPUP8* in cytokinin-related physiological functions rather than a primary role in flavin transport (Hu et al., 2023). Taken together, these findings support the notion that *AtPUP8* may have functionally specialized for substrates distinct from flavins, whereas *AtPUP5* has evolved a distinct specialization as a plasma membrane–localized protein contributing to flavin transport and localized riboflavin distribution in plants. Interestingly, some *AtPUP* members (e.g., *AtPUP7*, *AtPUP11*, and *AtPUP12*) appeared to negatively affect yeast growth under RF-supplemented conditions. One possible explanation is that expression of these transporters may perturb intracellular riboflavin homeostasis, potentially by affecting membrane transport processes. However, further investigation will be required to clarify the underlying mechanism.

Assessment of flavin coenzyme uptake in *AtPUP5*-overexpressing plants further showed that application of FMN or FAD did not result in detectable increases in cellular FMN or FAD levels, whereas RF levels were markedly elevated (Figure 4). In contrast, yeast-based assays showed that expression of *AtPUP5* was associated with increased intracellular accumulation of RF and FMN, whereas accumulation of FAD was substantially less efficient. The modest changes in intracellular FMN and FAD levels despite robust RF uptake suggest that imported flavins may be subject to rapid metabolic conversion and homeostatic control (Figure 2). Given their central metabolic roles, intracellular FMN and FAD levels are likely to be tightly regulated. Thus, the lack of detectable increases in FMN and FAD levels in planta likely reflects rapid metabolic turnover and/or stringent homeostatic regulation, resulting in no net expansion of total coenzyme pools. These observations are consistent with a role for AtPUP5 in riboflavin transport, whereas downstream flavin coenzyme levels are governed by metabolic buffering mechanisms. This distinction further supports a model in which plasma membrane–mediated riboflavin transport contributes to localized flavin distribution without directly perturbing global coenzyme homeostasis.

Despite the ability of *AtPUP5* overexpression to enhance RF accumulation, loss-of-function analyses revealed that *pup5*, *pup8*, and *pup5/8* mutants did not exhibit altered short-term or long-term RF uptake when assessed at the whole-plant level (Figures 5 and 6). Furthermore, long-term external RF treatment did not affect overall plant growth or development, including leaf area and flowering time, in either wild-type or mutant plants (Figure S5). Quantification of flavin levels in individual organs likewise revealed no significant differences in RF accumulation between wild-type and knockout lines when considering overall flavin accumulation across the plant (Figure 6). These findings indicate that *AtPUP5* and *AtPUP8* are not essential for bulk uptake of exogenously supplied RF at the whole-plant level under the conditions tested. Rather than serving as primary determinants of global flavin acquisition, *AtPUP5* appears to function in fine-tuning riboflavin distribution within specific tissues. Like cytokinin transport, which is mediated by multiple uptake and efflux transporters (Bürkle *et al*., 2003; Szydlowski *et al*., 2013), flavin uptake in plants may similarly involve a network of transporters that compensate for the loss of individual components. Such redundancy would allow maintenance of systemic flavin levels while permitting localized modulation of riboflavin distribution by specific transporters such as *AtPUP5*.

In contrast to the lack of effects on whole-plant RF uptake, loss of *AtPUP5* resulted in a pronounced and reproducible increase in RF levels in reproductive organs. RF accumulated to significantly higher levels in inflorescences of *pup5* plants than in wild-type plants, regardless of external RF treatment, and this phenotype was suppressed by complementation with *AtPUP5* (Figures 6 and 7). Moreover, RF levels were elevated in siliques and dry seeds of *pup5* (Figure 7), indicating that loss of *AtPUP5* affects flavin distribution at later stages of reproductive development. These observations suggest that *AtPUP5* may be involved in modulating riboflavin flux within reproductive tissues, thereby preventing excessive local accumulation. Rather than acting as a determinant of systemic flavin uptake, *AtPUP5* appears to function as a spatial regulator that fine-tunes riboflavin levels in specific developmental contexts. Such localized control may be particularly important during reproductive development, when metabolic demands and redox requirements are dynamically regulated. Because flavins function as essential redox cofactors for numerous oxidoreductases, localized regulation of riboflavin availability may contribute to maintaining appropriate redox balance during reproductive development.

Consistent with this interpretation, promoter–GUS analysis revealed strong *AtPUP5* expression in reproductive structures, including anthers, stigmas, and developing siliques (Figure 8). The spatial expression pattern of *AtPUP5* closely corresponds to the tissues in which RF overaccumulation was observed in *pup5* mutants, supporting a role for *AtPUP5* in spatial control of riboflavin homeostasis during reproductive development. This expression pattern is further supported by quantitative RT–PCR analysis, which confirms preferential expression of *AtPUP5* in reproductive tissues. Previous studies have shown that genes involved in flavin biosynthesis are transcriptionally regulated and respond to metabolic and environmental cues (Namba et al., 2024), whereas their spatial expression patterns in specific organs remain less well characterized. In this context, the preferential expression of *AtPUP5* in reproductive tissues suggests coordinated regulation of flavin metabolism and transport at the tissue level. Although the physiological functions of RF in reproductive organs of *Arabidopsis* remain largely unexplored, studies in *Capsicum pubescens* have reported RF accumulation in nectar, where it contributes to antimicrobial defense and attraction of pollinators (Magner et al., 2024). In addition, exogenous RF treatment has been reported to enhance seed stress tolerance in *rice* (Jiadkong et al., 2022; 2024). Together, these findings suggest that tight spatial regulation of riboflavin levels in reproductive tissues may have functional consequences for reproductive success and stress resistance. While the precise physiological roles remain to be elucidated, our results provide a molecular basis for such localized regulation.

Interestingly, while *AtPUP5* overexpression enhanced RF accumulation following external application, loss of *AtPUP5* led to RF overaccumulation in specific reproductive organs (Figures 3 and 7). This apparent dual effect–namely increased riboflavin accumulation in overexpression lines and organ-specific overaccumulation in loss-of-function mutants–is consistent with a model in which AtPUP5 functions as a membrane–localized protein associated with riboflavin transport that may facilitate riboflavin movement. In the absence of *AtPUP5*, altered riboflavin flux across cellular boundaries may contribute to its local accumulation within reproductive tissues. Such a phenotype is consistent with a model in which *AtPUP5* may facilitate riboflavin movement between adjacent tissues or cells, thereby preventing excessive local accumulation. This conceptual framework is summarized in Figure 10. Our results support a working model in which AtPUP5 contributes to the spatial regulation of riboflavin distribution in *Arabidopsis* reproductive tissues (Figure 10). AtPUP5 is localized to the plasma membrane and is preferentially expressed in inflorescences and developing siliques, consistent with a role in localized riboflavin homeostasis. In the absence of *AtPUP5*, riboflavin accumulates in reproductive organs, including seeds, suggesting that *AtPUP5* contributes to maintaining proper riboflavin distribution.

**Figure 10.**
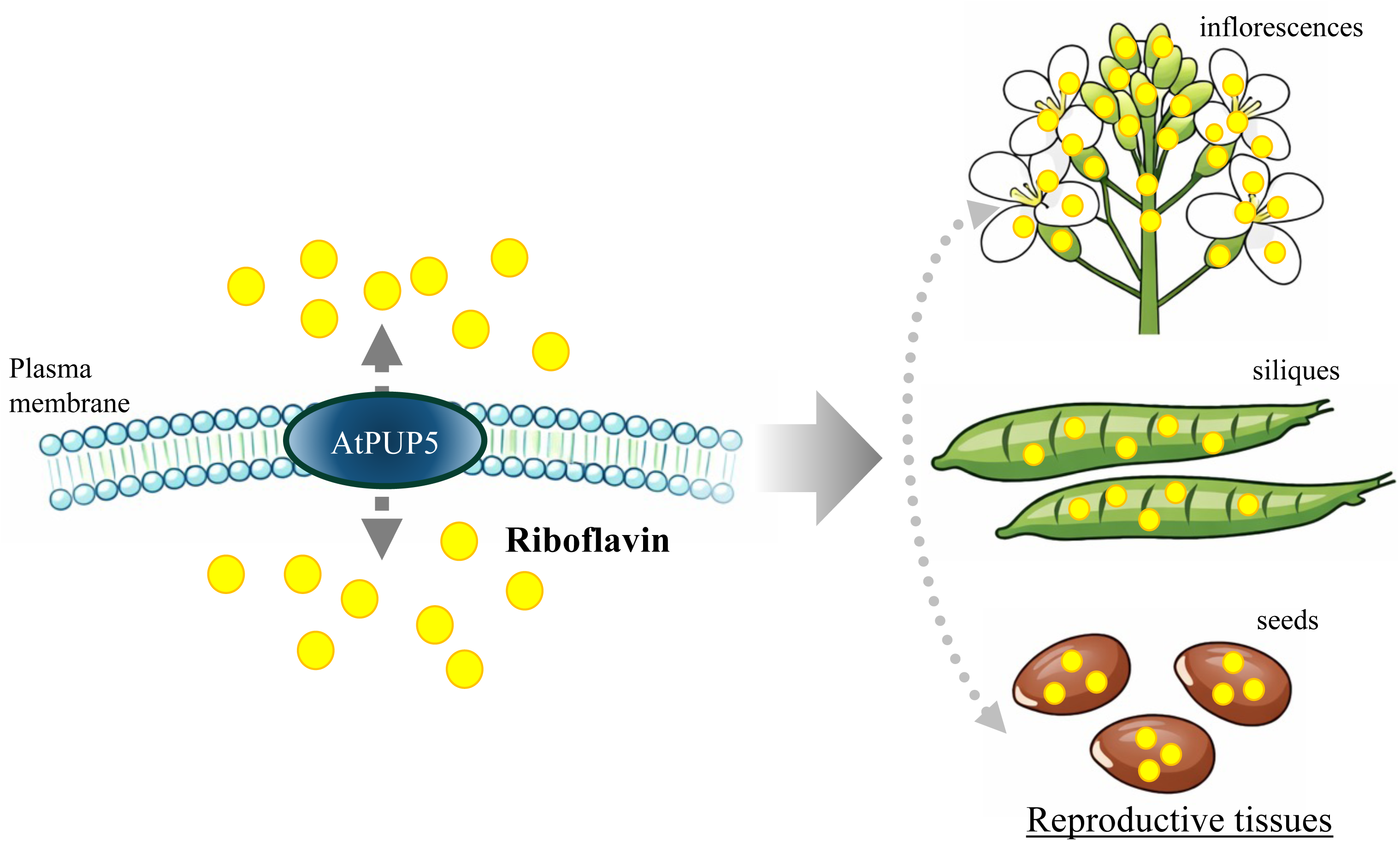
Proposed model for the role of AtPUP5 in the spatial regulation of riboflavin distribution in Arabidopsis reproductive tissues. AtPUP5 is localized to the plasma membrane and is associated with riboflavin distribution in reproductive organs, including inflorescences, siliques, and seeds. The model summarizes a conceptual framework based on the present study, in which AtPUP5 contributes to the spatial regulation of riboflavin distribution. The precise transport mechanism, substrate specificity, and directionality of AtPUP5 remain to be elucidated.

In summary, our study provides genetic and physiological evidence that the *Arabidopsis* purine permease *AtPUP5* contributes to riboflavin transport and spatial regulation of flavin distribution in plants. Using yeast complementation assays, transgenic overexpression lines, loss-of-function mutants, and promoter activity analyses, we demonstrate that *AtPUP5* is associated with riboflavin accumulation and plays an important role in regulating localized riboflavin distribution in reproductive tissues of *Arabidopsis*. Although our results support a role for *AtPUP5* in riboflavin transport, the precise transport mechanism remains to be clarified. Biochemical transport assays using purified membrane systems or heterologous expression platforms will be required to determine whether riboflavin is a direct substrate of *AtPUP5*, as well as to define its substrate range, transport directionality, and kinetic properties. *Arabidopsis* contains multiple PUP family members, raising the possibility that functional redundancy among related transporters contributes to maintaining systemic flavin homeostasis. In addition, other classes of membrane transporters may also participate in flavin transport, and such parallel transport systems could compensate for the loss of *AtPUP5*. Such redundancy may explain why loss of *AtPUP5* primarily affects riboflavin accumulation in specific reproductive organs without markedly altering global flavin levels at the whole-plant level. Together, our findings provide a molecular framework for plasma membrane–mediated control of riboflavin distribution in plants and highlight spatial regulation as an important component of flavin homeostasis during plant development.

## Materials and methods

### Plant materials and growth conditions

*Arabidopsis thaliana* ecotype Col-0 was used as a wild-type control. The T-DNA insertion mutants, *pup5* (CS868942) and *pup8* (SALK_137529) were obtained from the Arabidopsis Biological Resource Center. Seeds were sown in soil or on half-strength Murashige and Skoog (1/2 MS) medium containing 1% (w/v) sucrose. After three days of incubation at 4 °C in darkness, the seedlings were transferred to a growth chamber under standard conditions (16 h of light at 22 °C and 8 h of darkness at 20 °C), with a light intensity of 100 µmol photons m^-2^ s^-1^.

### Plasmid construction for AtPUP expression in yeast

The total RNA extracted from *Arabidopsis* aerial parts was reverse transcribed into first-strand cDNA using ReverTra Ace (Toyobo) and oligo (dT)_20_ primers. *AtPUP* cDNAs were amplified from the first-strand cDNA using the primer set provided in Table S1. The amplified cDNAs were inserted downstream of the galactose-inducible *GAL1* promoter of the multicopy vector pYES2 (Invitrogen) using In-Fusion Cloning technology (Takara).

### Generation of a yeast rib5 deletion mutant *(rib5Δ)* and yeast complementation assay

The *Saccharomyces cerevisiae* strain W303A (*MATa leu2-3,112 trp1-1 can1-100 ura3-1 ade2-1 his1-11,15*) was used in this study (Louis, 2016). Standard yeast culture media and genetic techniques were employed (Amberg and Burke, 2016). The media and growth conditions for the *rib5*Δ mutant and yeast complementation assays followed a previous report (Reihl and Stolz, 2005).

The *rib5*Δ mutant was generated by transformation of W303A with a *rib5*Δ::*kanMX6* disruption cassette amplified using the primer set listed in Table S1. The *rib5*Δ mutant was grown on YPDA medium (2% D-glucose, 2% peptone, 1% yeast extract, and 0.04% adenine) supplemented with G418 (200 mg L^-1^) and riboflavin (20 mg L^-1^). Each *AtPUP* cDNA-carrying pYES2 vector was transformed into the *rib5*Δ mutant using standard yeast transformation techniques (Kawai et al., 2010). Transformed *rib5*Δ mutant cells were plated on synthetic defined medium lacking uracil containing 2% D-glucose (SD/−Ura) supplemented with 20 mg L^-1^ riboflavin. The pYES2 vector lacking an *AtPUP* cDNA insert was used as a negative control.

For yeast complementation assays, cells were cultured for 24 h at 30 °C in 10 mL of SD/−Ura medium containing 20 mg L^-1^ riboflavin. Cells were then washed twice with sterile water, transferred to 10 mL of SD/−Ura without riboflavin, and cultured for 24 h to deplete intracellular riboflavin. After washing, the OD_600_ of each cell suspension was adjusted to 1.0. Yeast suspensions were serially diluted 10-, 10^2^-, or 10^3^-fold with sterile water and spotted onto SD/−Ura or synthetic defined medium minus uracil containing 2% D-galactose (SG/−Ura) agar plates supplemented with varying concentrations of riboflavin. Growth was recorded after 72 h of incubation at 30 °C.

### Flavin uptake experiments in yeast

The *S. cerevisiae rib5*Δ mutant cells carrying *AtPUP5*, *AtPUP8*, or the pYES2 empty vector were cultured for 24 h at 30 °C in 10 mL of SD/−Ura medium containing 50 μM riboflavin. After incubation, the OD_600_ was measured, and the yeast suspensions were diluted into 20 mL of SD/−Ura medium containing 50 μM riboflavin to achieve an OD_600_ of 0.1. The cultures were then incubated for an additional 24 h. Cells were collected by centrifugation, washed twice with sterile water, resuspended in 10 mL of SD/−Ura without riboflavin, and cultured for 24 h to deplete intracellular riboflavin. After washing, the cells were resuspended in 10 mL of SG/−Ura medium containing 50 μM riboflavin, flavin mononucleotide (FMN), or flavin adenine dinucleotide (FAD) and incubated for 12 h.

For short-term uptake experiments, yeast cells depleted of intracellular riboflavin were resuspended in SG/−Ura medium to induce *AtPUP* expression. After 6 h of incubation, each flavin compound was added to a final concentration of 50 μM, and yeast cells were collected at 2, 4, and 6 min post-addition. All collected yeast cells were washed three times with cold sterile water, harvested by centrifugation, frozen in liquid nitrogen, and stored at −80 °C until analysis.

### Plasmid construction and transformation of Arabidopsis

To construct the plasmid for constitutive overexpression of *AtPUP5*, we used a modified pRI201-AN vector (Takara, Kyoto, Japan) in which the *NPTII* gene was substituted with the *HPT* gene (Namba et al., 2024). The AtPUP5 cDNA was amplified from the first-strand cDNA and inserted downstream of the CaMV 35S promoter in the modified pRI201-AN vector (pRI201/35S-*AtPUP5*). For generating *AtPUP5* complemented *Arabidopsis* plants, the CaMV 35S promoter in pRI201/35S-*AtPUP5* was replaced with a 2,000-bp upstream region of the *AtPUP5* start codon (pRI201/*PUP5pro-AtPUP5*). The pGWB506 and pRI201-AN-GUS (Takara) vectors were used for expressing the GFP-AtPUP5 fusion protein and performing the AtPUP5 promoter GUS assay, respectively. The *AtPUP5* cDNA was cloned into pGWB506, allowing expression of the AtPUP5 protein fused to GFP at its N-terminus under the control of the CaMV 35S promoter. For the promoter GUS assay, the 2,000-bp upstream region of the *AtPUP5* translation start site was inserted upstream of the GUS gene. All plasmids were constructed using the primer sets listed in Table S1 and In-Fusion Cloning technology. *Agrobacterium tumefaciens* strain C58, transformed with the constructs *via* electroporation, was used to infect *Arabidopsis* wild-type plants (Col-0) or *pup5* mutants through the floral dip transformation method. T_1_ seedlings were selected on half-strength MS (1/2 MS) medium containing 1% (w/v) sucrose and 20 mg L^−1^ hygromycin for 2 weeks before being transferred to soil.

### Semi-quantitative and quantitative real-time PCR analysis

The total RNA extracted from *Arabidopsis* aerial parts was reverse transcribed into first-strand cDNA using ReverTra Ace (Toyobo) and oligo (dT)_20_ primers. In the semi-quantitative RT-PCR analysis, PCR amplification was conducted through 22–35 cycles of denaturation at 95 °C for 30 s, annealing at 55 °C for 30 s, and extension at 72 °C for 60 s, followed by a final extension step at 72 °C for 10 min. The uniform loading of each amplified cDNA was verified using the *ACTIN2* control PCR product. Quantitative real-time PCR analysis was carried out following the procedures outlined in a previous study (Namba et al., 2024). Primer sequences are provided in Table S1.

### Flavin uptake experiments in planta

The experimental system using cellophane sheets was established according to a previous report (Hachiya et al., 2021). A sterile cellophane sheet was placed on the solidified medium with sterilized tweezers using aseptic technique. Plants were grown on half-strength Murashige and Skoog (1/2 MS) solid medium containing 1% (w/v) sucrose on cellophane sheets for 2 weeks under standard growth conditions. Seedlings grown on cellophane sheets placed on solid medium were transferred together with the cellophane sheets to filter paper soaked in liquid 1/2 MS medium supplemented with or without 50 µM riboflavin (RF), flavin mononucleotide (FMN), or flavin adenine dinucleotide (FAD) and incubated in the dark to prevent photodegradation of flavins. After 1, 3, and 6 h of incubation, plants were separated into shoots and roots and harvested, thoroughly washed, frozen in liquid nitrogen, and stored at −80 °C until analysis.

### Long-term external riboflavin treatment and phenotyping of plants

All plants were cultivated in soil within a growth chamber under standard conditions and irrigated with water (Mock) or water supplemented with 50 µM riboflavin (RF). At 3 weeks after sowing, plants were photographed and the rosette area was measured using ImageJ software. Flowering time was defined as the day on which the inflorescence stem reached 1 cm in length. At that time, rosette leaves were counted to determine the flowering time by leaf number, excluding cauline leaves. At 40 days after sowing, all plants were separated into their respective organs and harvested 4 h after the onset of illumination, frozen in liquid nitrogen, and stored at −80 °C until analysis.

### Measurement of cellular flavins

Frozen organs of *Arabidopsis* plants (approximately 20–50 mg) were ground with zirconia beads (5.0 mm) using a multi-bead shocker (Yasui Kikai, Osaka, Japan) and homogenized in 0.5 mL of 50% methanol (v/v). For yeast cells, the frozen samples (approximately 20–50 mg) were ground with zirconia/silica beads (0.5 mm) in 0.2 mL of 50% methanol (v/v) using the same multi-bead shocker. The resulting extracts were heated at 80 °C for 10 min in darkness to prevent flavin degradation. After heating, the samples were centrifuged at 20,000 × g for 15 min at 4 °C. The supernatants were dried in darkness using a rotary evaporator and subsequently reconstituted in the mobile phase. These reconstituted extracts were filtered through a 0.2 µm filter (Millipore) before analysis. Flavin content was quantified using high-performance liquid chromatography (HPLC) with a COSMOSIL 5C18-MS-II column (4.6 × 250 mm, Nacalai Tesque, Kyoto, Japan). The mobile phase consisted of 10 mM NaH□PO□ (pH 5.5) containing 30% methanol (v/v) at a flow rate of 0.5 mL min□¹. Fluorescence detection was performed with an excitation wavelength of 445 nm and an emission wavelength of 530 nm. Quantification was performed by comparison with external standard curves generated using authentic riboflavin, flavin mononucleotide, and flavin adenine dinucleotide standards.

### Histochemical analysis of GUS reporter activity

Transgenic plants were immersed in ice-cold 90% acetone and fixed by standing on ice. After fixation, acetone was removed using 100 mM sodium phosphate buffer (pH 7.0). The plant samples were incubated in GUS staining solution containing 100 mM sodium phosphate (pH 7.0), 0.5 mM K4[Fe(CN)6], 0.5 mM K3[Fe(CN)6], 0.1% (v/v) Triton X-100, and 1 mM

5-bromo-4-chloro-3-indolyl-β-D-glucuronic acid (X-Gluc) for 2 h at 30 °C. A slight vacuum was applied before incubation to facilitate substrate infiltration. The staining solution was removed, and the tissue was washed three times with 70% (v/v) EtOH. It was then immersed in a decolorizing solution containing EtOH and acetic acid (6:1, v/v), and rotated overnight at 37 °C. The decolorizing solution was removed, and the tissue was washed three times with 70% (v/v) EtOH. It was then immersed in a clearing solution containing chloral hydrate: glycerol: water (8 g: 1 mL: 2 mL) and rotated for 2–3 h at room temperature to clear the tissue. Tissues were observed in the clearing solution using an all-in-one fluorescence microscope (KEYENCE BZ-X700).

### Transient expression of nGFP-AtPUP5 and subcellular localization analysis by particle bombardment

Subcellular localization of the AtPUP5 protein was analyzed by transient expression in *Arabidopsis thaliana* leaf epidermal cells using particle bombardment. Fully expanded rosette leaves were collected from 3–5-week-old plants grown under standard growth conditions. Detached leaves were placed with the adaxial side facing up on solid Murashige and Skoog (MS) medium and equilibrated at room temperature prior to bombardment.

For co-localization analysis, plasmids encoding an N-terminal GFP fusion of *AtPUP5* (nGFP–AtPUP5; pGWB506/AtPUP5) and a plasma membrane–localized RFP marker (AtSYP132–RFP; pB5tdGW/AtSYP132) (Nishimura et al., 2016) were co-introduced into leaf epidermal cells. Tungsten particles (approximately 1.0 µm in diameter) were coated with plasmid DNA using CaCl_2_–spermidine–mediated precipitation. Briefly, plasmid DNA was mixed with tungsten particles, followed by sequential addition of CaCl_2_ and spermidine under continuous vortexing to allow DNA precipitation onto the particles. The DNA-coated particles were washed with absolute ethanol, resuspended in ethanol, and loaded onto macrocarriers according to the manufacturer’s instructions.

Particle bombardment was performed using a helium-driven particle delivery system under vacuum (TANAKA GIE-III IDELA). Leaves were bombarded at a target distance and helium pressure optimized for *Arabidopsis* leaf tissue. Following bombardment, samples were incubated on MS medium at approximately 22 °C under dark conditions overnight to allow transient expression of the introduced constructs. Fluorescence signals were observed using a confocal laser scanning microscope (Leica TCS SP5).

### Statistical analyses

Statistical analyses were performed using Student’s *t*-test. All calculations were performed using at least three independent biological replicates.

## Acknowledgment

We are grateful to Prof. Tsuyoshi Nakagawa (Shimane University) for providing the pGWB6 vector, to Asst. Pro. Takashi Akihiro (Shimane University) for the pYES2/OsPUP5 construct, and to Assoc. Prof. Koji Nishimura (Shimane University) for the pB5tdGW/AtSYP132 vector. We also thank Prof. Yasuhiro Matsuo (Shimane University) for providing the *Saccharomyces cerevisiae* strains and for his technical support and valuable advice on yeast experiments. We are sincerely grateful to Prof. Takanori Maruta (Shimane University) for insightful discussions and valuable advice throughout this study. We acknowledge Ayaka Daihuku and Fumina Nagai for their assistance and support in this research.

## Author Contributions

T.O. conceived and designed the research. R.S. and H.K. performed most of the experiments. T.S. conducted the yeast screening and intracellular localization analyses, and M.K. generated the yeast mutants. T.I. and T.O. supervised the research and contributed to data interpretation. R.S., H.K., and T.O. analyzed the data. T.O. wrote the manuscript with input from all authors. All authors read and approved the final manuscript.

## Conflict of Interest

The authors declare no conflict of interest.

## Funding

This work was supported by JSPS KAKENHI Grant Numbers JP18K05439, JP21K05382, and JP26K08631 (to TO).

## Data Availability

The data supporting the findings of this study are available within the article and its Supplementary information. Raw data underlying the figures are available from the corresponding author upon reasonable request.

## Abbreviations

FAD: flavin adenine dinucleotide
FMN: flavin mononucleotide
RF: riboflavin
PUP: purine permease

## Notes

### Competing Interest Statement

The authors have declared no competing interest.

### Summary of Updates

We have (i) improved the clarity of Figure S5, (ii) included a new schematic model (Figure 10) summarizing the main findings of the study, (iii) added quantitative RT-PCR analysis data to support the tissue-specific expression pattern of AtPUP5 (Figure S7), and (iv) expanded the Discussion to better integrate our findings with previous studies on flavin metabolism and gene expression patterns.

