## Supplemental Figures for "Purine permease 5 contributes to riboflavin distribution in *Arabidopsis* reproductive organs"

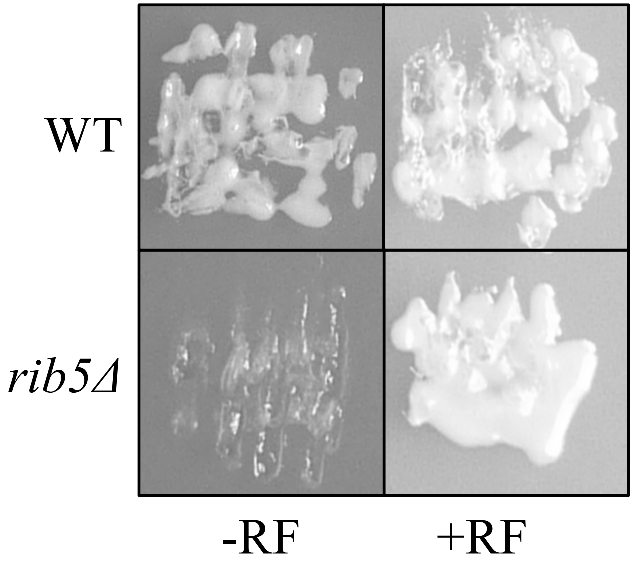


**Figure S1 Growth of the *Saccharomyces cerevisiae rib5Δ* mutant requires supplementation with high concentrations of RF**
*S. cerevisiae* wild-type (W303A) and *rib5Δ* mutant cells were plated on synthetic defined medium lacking uracil and containing 2% (w/v) D-glucose (SD/−Ura), supplemented with or without riboflavin (20 mg $L^{-1}$), and were incubated at 30 ° C for 72 h.

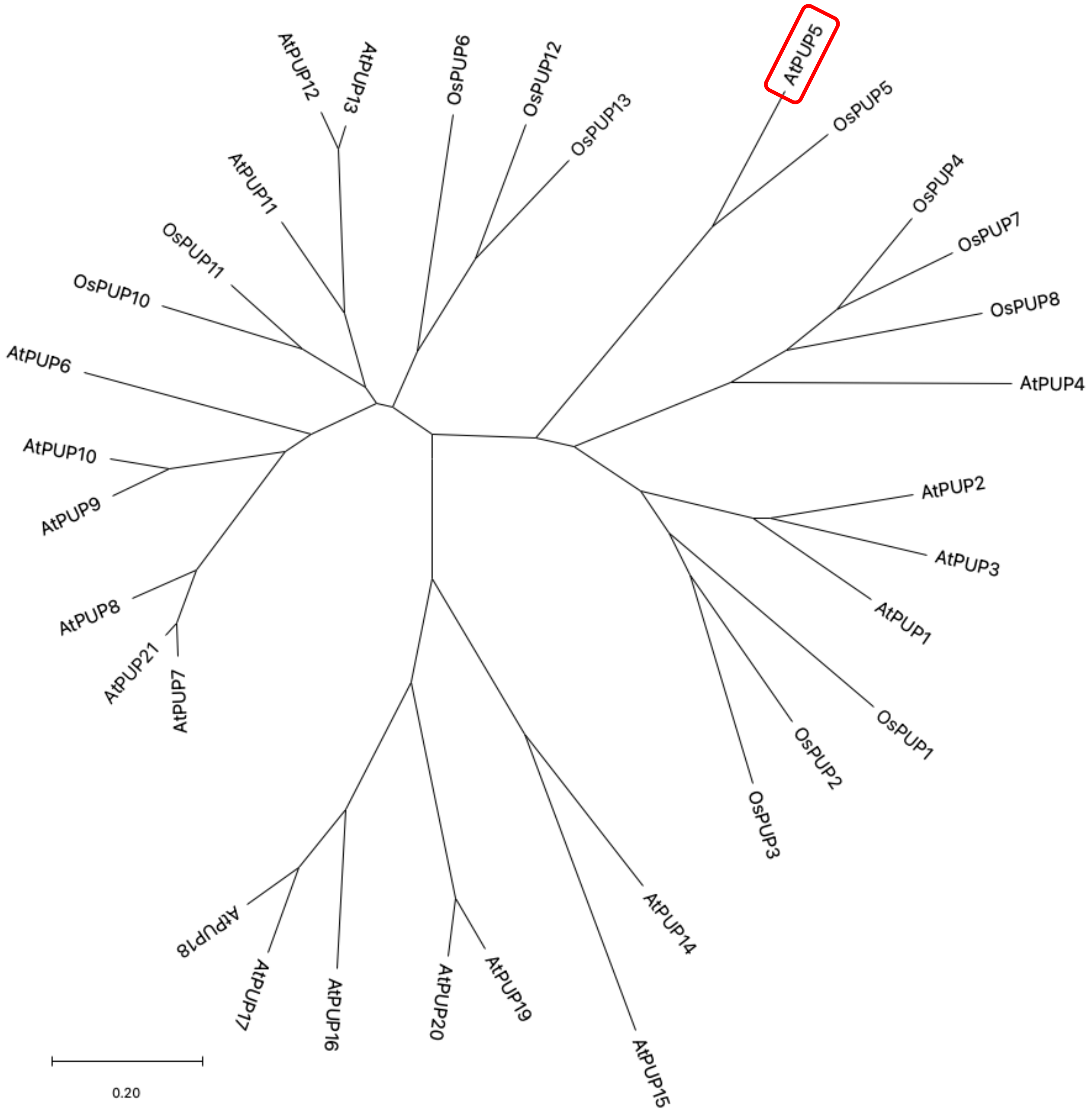


**Figure S2 Phylogenetic analysis of purine permease (PUP) proteins from *Arabidopsis thaliana* and *Oryza sativa***

Amino-acid sequences of purine permease proteins from *Arabidopsis thaliana* and *Oryza sativa* were aligned using MUSCLE implemented in MEGA 11. The phylogenetic tree was constructed using the Neighbor–Joining method with Poisson correction. Branch support was assessed by bootstrap analysis with 1,000 replicates .

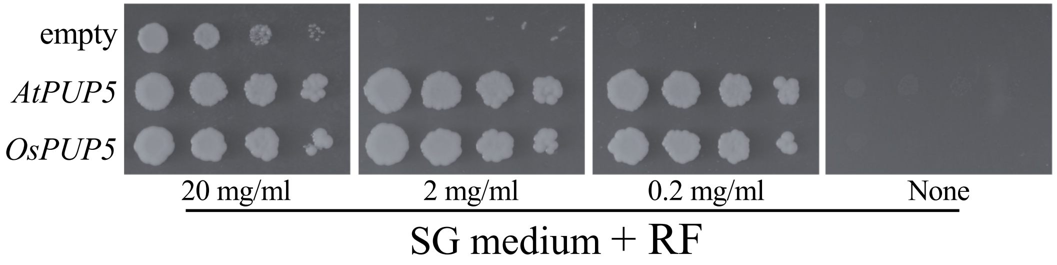


**Figure S3 Functional complementation of the *Saccharomyces cerevisiae rib5Δ* mutant by *AtPUP5* and its rice homolog *OsPUP5***
Complementation assays were performed using a *rib5Δ* yeast mutant transformed with an empty pYES2 vector (negative control) or pYES2 constructs harboring *Arabidopsis thaliana AtPUP5* or *Oryza sativa OsPUP5*. Transformed strains were spotted onto synthetic defined medium lacking uracil supplemented with riboflavin and incubated under the same conditions as described in Figure 1.

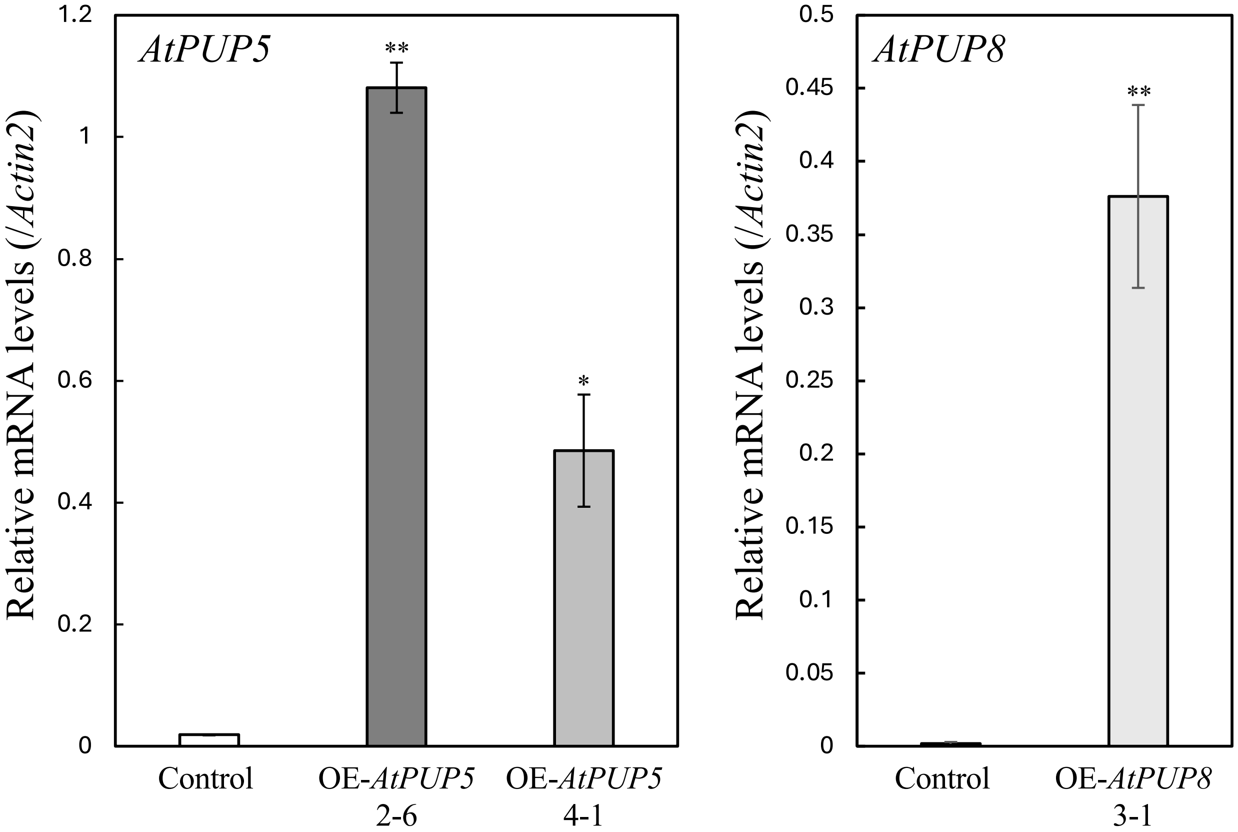


**Figure S4 Transcript levels of *AtPUP5* and *AtPUP8* in transgenic *Arabidopsis* plants constitutively overexpressing *AtPUP5 or AtPUP8***
Control plants (wild-type plants transformed with the empty vector), two independent *AtPUP5*-overexpressing lines, and one independent *AtPUP8*-overexpressing line were analyzed. Plants were grown on half-strength Murashige and Skoog (1/2 MS) medium supplemented with 1% (w/v) sucrose for 2 weeks under standard growth conditions, and transcript levels were quantified by quantitative real-time PCR. Data are presented as means ± SE from three biological replicates (n = 3). Asterisks indicate statistically significant differences compared with the empty vector control (Student's *t*-test; $^{*}P < 0.05$ and $^{**}P < 0.01$).

(a)

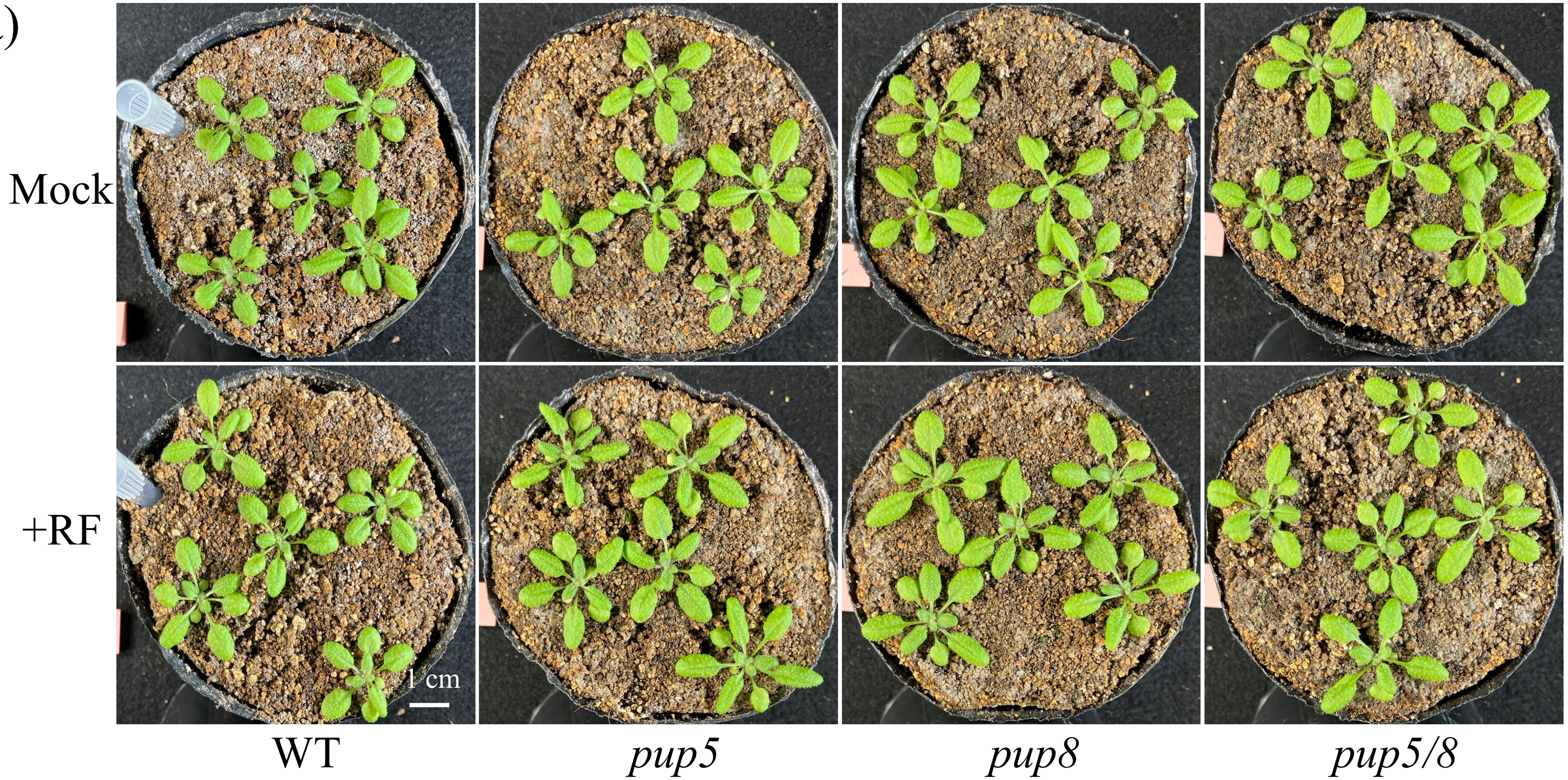


Figure S5-1

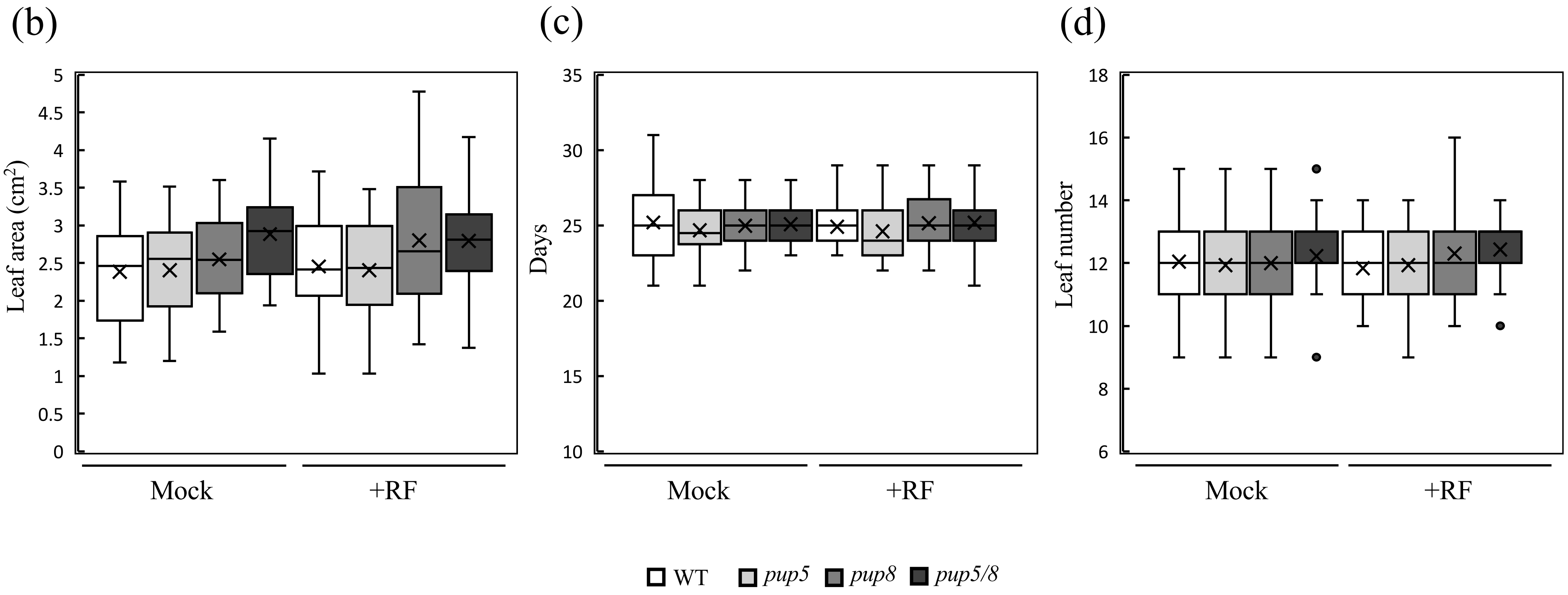


**Figure S5 Phenotypic characterization of the *pup5, pup8, and pup5/8* mutant plants grown in soil following long-term riboflavin treatment**

Wild-type plants and *AtPUP* knockout mutants (*pup5*, *pup8*, and *pup5/8*) were grown in soil for 40 days under standard growth conditions and irrigated with water (Mock) or water supplemented with 50 μM riboflavin (RF). (a) Representative photographs and (b) leaf area of wild-type and pup mutant plants grown for 3 weeks after sowing. For leaf area measurements, n = 30 plants per genotype; the experiment was conducted twice with similar results. Flowering time was evaluated by (c) days to bolting and (d) leaf number at bolting in wild-type and pup mutant plants (n = 43–45); the experiment was conducted twice with similar results. The lines within the boxes are the medians, and the lower and upper hinges represent the first and third quartiles. Cross marks indicate the mean values.

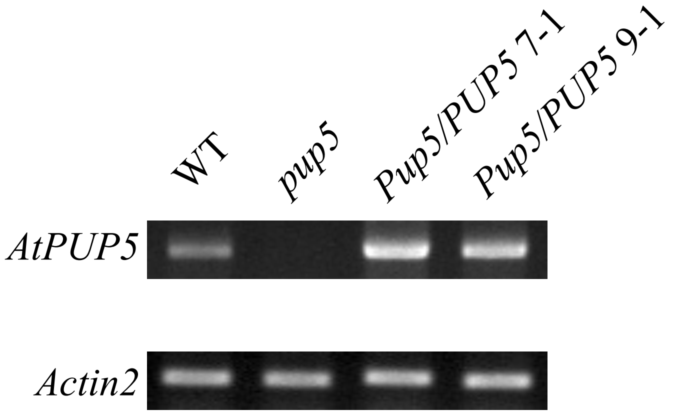


**Figure S6 Semi-quantitative RT–PCR analysis of *AtPUP5* expression in *pup5* mutant and complemented lines**
Wild-type plants, the *pup5* mutant, and two independent *AtPUP5*-complemented lines (*pup5*/*PUP5*) were analyzed. Plants were grown on half-strength Murashige and Skoog (1/2 MS) medium supplemented with 1% (w/v) sucrose for 2 weeks under standard growth conditions, and *AtPUP5* transcript levels were examined by semi-quantitative RT–PCR.

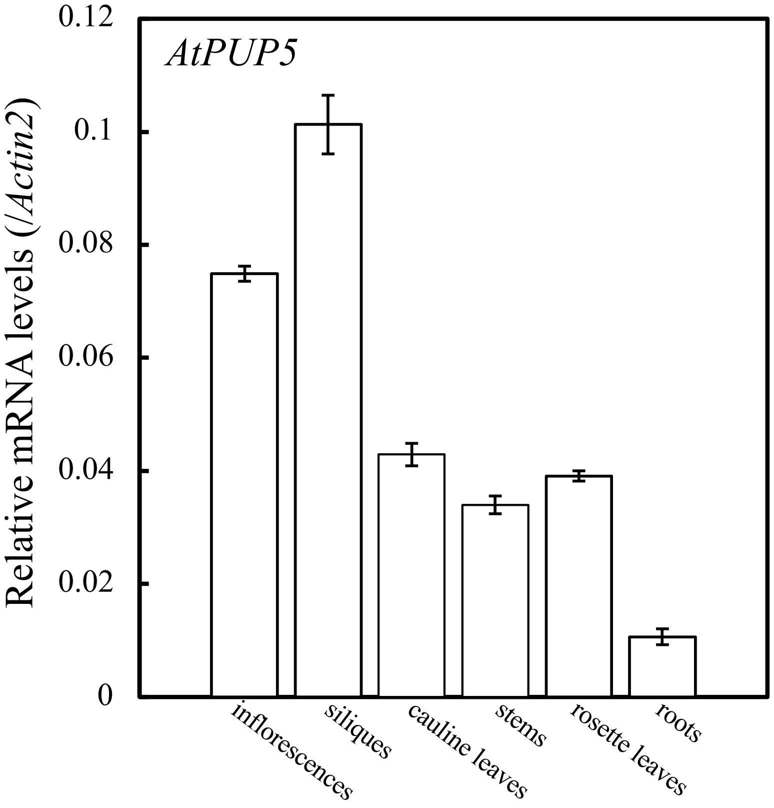


**Figure S7 Tissue-specific expression of *AtPUP5* analyzed by quantitative RT–PCR.**
Wild-type plants were grown in soil for 40 days under standard growth conditions. At 40 days, organs from wild-type plants were harvested 4 h after the onset of illumination, and *AtPUP5* transcript levels were quantified by quantitative real-time PCR. Data are presented as means ± SE (n = 3).
